# Increased birth rank of homosexual males: disentangling the older brother effect and sexual antagonism hypothesis

**DOI:** 10.1101/2022.02.22.481477

**Authors:** Michel Raymond, Daniel Turek, Valerie Durand, Sarah Nila, Bambang Suryobroto, Julien Vadez, Julien Barthes, Menelaos Apostoulou, Pierre-André Crochet

**Author notes:** Corresponding author : Michel RAYMOND, Institute of Evolutionary Sciences, CC065, University of Montpellier, 34095 Montpellier Cedex 05, France.

## Abstract

Male homosexual orientation remains a Darwinian paradox, as there is no consensus on its evolutionary (ultimate) determinants. One intriguing feature of homosexual men is their higher male birth rank compared to heterosexual men. This can be explained by two non-exclusive mechanisms: an antagonistic effect (AE), implying that more fertile women have a higher chance of having a homosexual son and to produce children with a higher mean birth rank, or a fraternal birth effect (FBOE), where each additional older brother increases the chances for a male embryo to develop a homosexual orientation due to an immunoreactivity process. However, there is no consensus on whether both FBOE and AE are present in human populations, or if only one of these mechanisms is at play with its effect mimicking the signature of the other mechanism. An additional sororal birth order effect (SBOE) has also recently been proposed. To clarify this situation, we developed theoretical and statistical tools to study FBOE and AE independently or in combination, taking into account all known sampling biases. These tools were applied on new individual data, and on various available published data (two individual datasets, and all relevant aggregated data). Support for FBOE was apparent in aggregated data, with the FBOE increasing linearly with fertility. The FBOE was also supported in two individual datasets. An SBOE is generated when sampling in presence of FBOE, suggesting that controlling for FBOE is required to avoid artefactual SBOE. AE was not supported in individual datasets, including the analysis of the extended maternal family. The evolutionary implications of these findings are discussed.

## INTRODUCTION

Male homosexual orientation, i.e. preferential attraction of male subjects to same-sex partners for sexual intercourse and/or romantic relationships, is an evolutionary enigma. This is because preference for male-male relationships is partially heritable (Bailey et al., 2000; Långström et al., 2010), and is associated with a fertility cost with a 70-100% decrease in offspring number (Iemmola and Camperio-Ciani, 2009; Nila et al., 2018; Rieger et al., 2012; Vasey et al., 2007) Also, male homosexual orientation is surprisingly common in many societies (2%–6% in Western countries) for such a costly trait (Apostolou, 2020; Berman, 2003). The origin of male homosexual orientation has long been a matter of interest, and several evolutionary hypotheses have been proposed, mostly involving kin selection or antagonistic pleiotropy (see Barthes et al., 2015, Gavrilets and Rice, 2006, and Apostolou, 2020 for reviews).

In this paper, we will examine some of the empirical evidence linked to one of these evolutionary hypotheses, the antagonistic pleiotropy hypothesis. This hypothesis explains the persistence of male homosexuality by a sex-antagonistic effect (AE). Several studies have indeed reported differences in fertility between families of homosexual and heterosexual men (Camperio-Ciani et al., 2009, 2004a; Iemmola and Camperio-Ciani, 2009), usually reported as a Female Fecundity Effect (FFE), and the authors proposed that families of homosexual men display a higher fecundity of female relatives from the maternal side (e.g. maternal aunts), compared with the rest of the population, in accordance with the AE hypothesis.

Another line of research has attempted to decipher the proximal determinants of male homosexual orientation. Local environmental effects within the family for the sexual orientation of men has been suspected for nearly a century, and two main directions have been explored (for a short review, see Wampold (2018)). The first focused on birth order, and concluded that homosexual men had more older siblings than heterosexual men (e.g. Slater, 1962); the second focused on sex-ratio, and concluded that homosexual men had more brothers than sisters (e.g. Lang, 1940). These two observations were reconciliated when it was proposed that homosexual men had more older brothers than heterosexual men (Blanchard and Bogaert, 1996a). This finding, commonly referred to as the fraternal birth order effect (FBOE), has been found repeatedly independently in Western (e.g., Blanchard, 2018a, 2018b; Blanchard and Bogaert, 2004; Bogaert and Skorska, 2011) and non-Western countries such as Turkey, Iran, Hong Kong, Samoa, and Indonesia (Blanchard, 2018c; Li and Wong, 2018; Nila et al., 2019). The underlying mechanism of the FBOE is proposed to be biological and prenatal, since homosexual orientation is influenced neither by the number of non-biological older brothers nor by the amount of time spent with biological or non-biological older brothers (Bogaert, 2006). The proposed explanation is a maternal immune reaction to successive male pregnancies, with each male foetus increasing the likelihood of an immune response from the mother. This maternal immune reaction would lead to an alteration of the typical development of sexually dimorphic brain structures relevant to the sexual orientation of the foetus (Bogaert and Skorska, 2011). Recently, possible molecular evidence of this specific immune reaction has been presented (Bogaert et al., 2018). Yet, these two lines of research, one on an evolutionary explanation (AE), and another on a proximate explanation (FBOE), leave many open questions.

Firstly, it is still unclear whether the FBOE is universal. The FBOE is not always found, even in some large samples from UK, Canada, or Australia (Bogaert, 1998; Kishida and Rahman, 2015; Rahman et al., 2008; Zietsch et al., 2012, but see Blanchard and VanderLaan, 2015). It is thus possible that the FBOE only operates in some populations. Alternatively, the FBOE could be restricted to subcategories of homosexual men (i.e. FBOE leads to only certain subcategories of male homosexuality), as suggested by Swift-Gallant et al. (2018). Additionally, the FBOE is sometimes described from samples which are not comparable. For example, several meta-analyses (Blanchard, 2018c, 2018a, 2018b; Blanchard et al., 2021, 2020) testing for an FBOE in homosexual men across multiple studies include data from transexuals, pedophiles, hebephiles, or gender-dysphoria individuals. Similarly, other studies have focused on specific individual categories, such as sex offenders, psychoanalytic patients, individuals treated with feminizing hormones, clinically obsessional patients, or patients with paraphilic behaviours such as masochism, fetishism, and transvestism (e.g. Blanchard et al., 2012, 1998). As these different situations are drawn from highly non-representative populations (Zietsch, 2018), and are not necessarily the result of similar determinants as those for homosexuality, or could represent extreme values from a continuum, considering them could introduce some biases.

Secondly, a sororal birth order effect (SBOE) acting alongside the FBOE has been described several times, e.g. in UK (King et al., 2005), Finland (Kangassalo et al., 2011), Samoa (VanderLaan and Vasey, 2011; Vasey and VanderLaan, 2007), Canada (Swift-Gallant et al., 2018), Netherlands (Ablaza et al., 2022), or in participants of a BBC internet survey (Blanchard and Lippa 2021). Based on these findings, a recent meta-analysis proposed the presence of a pervasive SBOE, in addition to the FBOE (Blanchard et al., 2021), even if this SBOE is generally not as strong as the FBOE. A further complication arises from the fact that an SBOE could in theory be a side-effect of an FBOE: if the sex ratio is even, sampling individuals with more older brothers also means sampling individuals with more older sisters. If this were the case, homosexual men would generally have more older sisters than heterosexual men even if the only causal effect on the probability of being homosexual is the number of older brothers. It is thus unclear if explanations for the SBOE should be sought independently or not from this additional sibling effect.

A third --and quite problematic-- question is whether the FBOE and AE are both at play in human populations. Indeed, a higher fertility of mothers of homosexual men implies that, when sampling homosexuals from a population, the mean birth order of homosexuals will be higher, on average, than the mean birth order of heterosexuals. Conversely, if fertility varies within a population independent of the occurrence of homosexuality, sampling high birth ranks (as is the case when sampling homosexuals in the presence of the FBOE) will generate a sample from high-fertility mothers. The two phenomena (FBOE and AE) thus lead to similar predictions of a higher birth rank of homosexuals and a higher fertility of families of homosexuals from population samples, and thus cannot be easily distinguished even if they rely on very different mechanisms: a plastic effect (maternal effect) in the case of FBOE and a genetic effect in the case of the AE. This problem of causal attribution has been previously identified (e.g. VanderLaan and Vasey, 2011; Zietsch et al., 2008), and formally exposed (Khovanova, 2020), and three main methods have been proposed to study FBOE while controlling for variation in female fecundity.

This first is a statistical control of fecundity: raw number of older brothers are not transformed, but family size is used as a control variable, for example as a dependent variable in a regression (e.g. Ablaza et al., 2022; Nila et al., 2019). Second, transforming the raw data using various various metrics controlling for family size: the general form of these metrics is (*X + a*)/(*N + b*), where *X* is the number of older brothers (or any other sibling category under study), *N* is the total number of siblings, and {*a, b*} are two scalars. Values of these scalars vary according to authors: {-*1, −1*} for Slater (1962), {*½, 1*} for Berglin (1980), and {*¼, 1*} or {*⅓, −X+1*} for Blanchard (2014). These metrics have some drawbacks, e.g. Slater’s index is not defined for only-children (*N* = 1), see Blanchard (2014) for further comments on these metrics. Other metrics have thus been subsequently proposed, based on ratio between the odds of observing an older brother for homosexuals, and the same odds for heterosexuals (OBOR, Blanchard, 2018c, 2018b), or based on the ratio of older brothers to older sisters, relative to the same ratio for heterosexuals (OR, Vilsmeier et al., 2021). Third, a data restriction: only families with a fixed number of children (i.e. 1 or 2) are considered (Khovanova, 2020). This can lead, in populations displaying a relatively high fecundity, to discarding more than 60 % of the sampled individuals (Blanchard et al., 2020).

Results using these methods indicated that FBOE was unlikely to be a sampling artefact, at least in some populations, although there is no consensus at a larger scale: several meta-analyses using an odds-ratio metric concluded that FBOE is generally present (Blanchard, 2018c, 2018c, 2018b), while a meta-analysis using another odds-ratio metric concluded that “*almost no variation in the number of older brothers in men is attributable to sexual orientation*” (Vilsmeier et al., 2021, but see Blanchard and Skorska, 2022). However, odds-ratios, and more generally all ratios, cannot be considered as reliable statistics for studying FBOE, even for non-restricted data, as this approach assumes that samples of heterosexuals and homosexuals are adequately matched for potentially confounding variables affecting sibship size, such as age or social economic status, which is not always the case (see Price and Hare, 1969). Generally, analysing ratios is rife with problems and should be avoided whenever possible (see e.g. Curran-Everett, 2013).

To study female fecundity without the interference of birth rank, one possibility is to use additional family data: the fecundity of maternal aunts, for example, is considered to be independent of the birth rank of the sampled individuals (Camperio-Ciani et al., 2004a; Iemmola and Camperio-Ciani, 2009; King et al., 2005). However, this is probably not the case, due to the correlation between the fecundity of the mother and her sisters (e.g. Anderton et al., 1987; Berent, 1953). Thus, even if an FBOE were acting alone (i.e. no AE), homosexuals will be sampled from families displaying a higher fecundity, and their maternal aunts are likely to display a higher fecundity as well due to this correlation (Zietsch et al., 2008). Another possible approach to control for birth rank is to consider only individuals with a specific birth rank, e.g. only first born or only second born (e.g. Blanchard et al., 2020; Ciani and Pellizzari, 2012; Khovanova, 2020), but this results in a significant reduction of the available data and hence of inferential power.

Results using these methods are ambiguous: the presence of AE is claimed when additional family data are considered, such as maternal aunts (e.g. Camperio-Ciani et al., 2004; Iemmola and Camperio-Ciani, 2009), but results are not controlled for birth rank. When only firstborn are considered, no evidence for AE was found (Blanchard, 2012), although the opposite conclusion was reached by Rieger et al. (2012). In a meta-analysis, when data are restricted to families with only one or only two sons, no AE is found (Blanchard et al., 2020).

To sum up, there is still no consensus on whether both FBOE and AE are present in human populations, or if only one of these mechanisms exists and its effect mimics the signature of the other mechanism when examining population samples. If both mechanisms act together, we also need to estimate the relative contribution of each for the higher birth rank of homosexual men and higher fertility of their families. Last, we do not know if the SBOE observed in several studies indicates a causal link between the number of older sisters and the probability of being homosexual, or if it is a by-product of the FBOE. This is a typical collinearity issue that we tried to address through various statistical approaches. To implement these approaches, we also had to first clarify some methodological issues with demographic parameter estimates derived from empirical sampling of human populations. Thus, we addressed the following points.

First, we derived a population-level relationship between mean birth rank and mean fertility in a random population sample, lacking FBOE, SBOE, or AE. This analytical relationship links the mean birth rank of individuals with the mean fertility of their population sample, allowing us to quantitatively estimate the role of FBOE, independently of any fertility effect, when mean birth rank deviates from its value predicted by the population fertility estimate. We also determined analytically and through simulations the expected number of older sisters as a function of the number of older brothers to test if the FBOE can indeed generate an SBOE.

Second, we checked if population samples from heterosexual and homosexual men gathered from the available literature followed the general expectation of mean birth rank given population fertility. If population samples of homosexuals deviated from this general expectation having higher birth rank than expected given the mean population fertility, this would support an FBOE effect.

Finally, we used two statistical frameworks to separate quantitatively the FBOE and AE effects in family data from individual homosexual and heterosexual men sampled in France, Indonesia and Greece. The first framework was “classical” linear modelling (here generalized linear mixed model or GLMM) to estimate the fertility of the women on the mother’s side (mothers and maternal aunts) after controlling for the birth rank of the focal subject (effectively comparing this familial fertility for first born, second born, etc.), allowing to test for AE after controlling for the FBOE. If, for a given birth rank, homosexuals had more sibling or cousins than heterosexuals, the AE hypothesis would be supported. The second framework was to use Bayesian inference implemented as a hierarchical model in NIMBLE (de Valpine et al., 2017) to test whether the effects of the FBOE, AE or both could be inferred from individual family data. NIMBLE generates simulated data based on different scenarios and provides quantitative support of the various scenarios from the empirical data. We evaluated several scenarios including solely or simultaneously the FBOE and the AE favouring male homosexuality and female fecundity. The FBOE was modelled after determining the best-fit function from the empirical data.

These different lines of reasoning allowed us for the first time to disentangle the relative explanatory power of birth rank (the FBOE) and of antagonist genetic factors (the AE) affecting fertility on patterns of sexual orientation simultaneously in aggregated data and in three independent data sets, and to provide several tools that will be useful for addressing these questions in other human populations.

## MATERIAL and METHODS

### Methodological developments

Analytical formulae were derived for some demographic variables, in absence of FBOE or AE, in order to generate null hypotheses for the analyses. This concerned the relationships between mean birth rank and mean fertility (Appendix 1), estimates of fecundity from sibling data (Appendix 2), and the sampling distribution of number of older brothers or older sisters (Appendices 1 and 1b). Simulation was further used to evaluate whether AE or FBOE generated a different relationship between birth rank and population fecundity for homosexual individuals than for the population as a whole. A total of 6000 families was generated, with fecundity drawn from a Poisson distribution of parameter λ, and with a 1:1 sex-ratio. The birth order of each sibling was recorded (male birth order among brothers, or female birth order among sisters). Sexual orientation for firstborn males (or for each sibling in absence of FBOE) was drawn from a binomial distribution of parameter *p,* with p = p_0_ = 0.05. AE was modelled as an increase in *λ* by a factor (*1 + β*), conjointly with an increase in *p_0_* by a factor (1+ α) (with α and β ≥ 0). FBOE was modelled using functions f5, f7, or f7’ of Table S1. From a random sample of 200 homosexual individuals, the mean birth order (MBO) and the mean fertility (MF) were calculated. This was replicated 50 times for a given λ, and this process was repeated for λ values from 2 to 8. The slope of the regression line MBO ∼ MF was calculated, and tested for a higher value than 0.5 by adding an appropriate offset term in the linear predictor.

### Aggregated family data

To find primary data on sexual orientation and family composition, we proceeded in two ways. First, we performed literature searches on accessible databases to find recent publications. Second, to find older data, we scanned cited literature. In addition, inspection of review articles ensured that no major older papers were overlooked (e.g. Blanchard, 2018b, 2018c, 2004, 1997; Blanchard et al., 2020a, 2020b, 2001; Bogaert and Skorska, 2011). Papers that present at least two samples of men, one homosexual and one heterosexual, along with the following information, were retained: number of individuals sampled, and the total number of each individual’s older brothers, older sisters and siblings. Data used in several papers were retained only once, e.g. Rahman et al. (2008) used in Kishida and Rahman (2015). When some required information was not found in the publication, we attempted to contact the authors to obtain the missing information. When the size of a sibling category was given only as a proportion this proportion was multiplied by the corresponding number of individuals and rounded to the nearest whole number to obtain the expected number for this sibling category. When further computation was required, the sample was not considered. In order to focus conservatively on typical homosexual or heterosexual individuals, data concerning sex-offenders, pedophiles, hebephiles, gender-dysphoria (i.e. conflict between gender identity and sex assigned at birth) individuals, psychoanalytic patients, individuals treated with feminizing hormones, hospital patients, clinically obsessional patients, patients with paraphilic behaviours like masochism, fetishism, and transvestism, or transexuals, were not considered. We also excluded samples concerned with children or adolescents (as the number of younger siblings may not be final), adopted individuals (as the biological sibling composition is generally not available) or twins (birth order is ambiguous). The French sample collected for individual data (see below) was also considered here in its aggregated form. Bisexual individuals were pooled with homosexuals, and pairs of samples with at least 50 individuals for each sexual orientation were further retained. For each sample, the mean number of older brothers was computed as *OB/N*, where *OB* is the total number of older brothers, and *N* is the number of sampled individuals. Then, the mean birth rank of men with respect to only their brothers was computed as *OB/N + 1*. Similarly, the mean birth rank of men with respect to only their sisters was computed as *OS/N +1*, where *OS* is the total number of older sisters. The mean fecundity was estimated as *Sibs/N*, where *Sibs* is the total number of older and younger siblings. The mean fecundity for only males or only females was computed as half this overall fecundity.

### Individual family data

Sampling in France was performed from August 2006 until July 2016 in public areas, research institutes, and within social networks (mainly in the cities of Montpellier and Paris). A targeted sampling procedure was performed: when a beach mostly frequented by individuals with a homosexual preference was sampled, a nearby beach with no particular attendance bias was also sampled. Friends and acquaintances of individuals reporting one or the other sexual preference were sampled. Upon agreement, a document describing the general purpose of the study and providing contact details of the person in charge (M.R.) was given to each participant. This document explicitly states that personal data will only be used for research purposes and that only global results –not individual data– will be published. Written informed consent was obtained from all participants. The protocols used to recruit individuals and to collect data were approved the French National Committee of Information and Liberty (CNIL) through the CNRS (approval #1226659). Each individual was privately and anonymously interviewed and was asked to report his date of birth, his self-declared sexual preference, the sex and birth order of each of his full and half siblings on the maternal side, the country of birth of his four grand-parents, the number of maternal aunts and corresponding number of cousins. Other personal and familial information was collected and will be analysed elsewhere. Individuals below 18 years of age (age of legal adulthood in France) were not considered. To reduce cultural heterogeneity, individuals with one or more grand-parent(s) born outside Europe were not further considered. Usable data were obtained from 512 men (Table 1). Eighteen of these men declared a bisexual orientation and were grouped in the homosexual preference category (removing them did not change qualitatively the results), giving a total of 271 men with a homosexual preference (52.9%), and 241 men with a heterosexual preference (47.1%). Mean (+/− SE) age was 33.8 (+/− 0.5) years (range 18.3 – 75.4), with the homosexual preference group being slightly younger than the heterosexual group: 32.0 (+/− 0.6) vs 35.8 (+/− 0.9) years, Wilcoxon test, W = 36838, *P* = 0.012.

**Table 1.** Aggregated data collected from published studies. For each pair of homosexual and heterosexual male samples, the number of focal individuals (N), their total number of older brothers (OB), older sisters (OS), and siblings (older and younger) are shown, as well as the origin of the sample and the reference.

| Reference | Sample origin | Sexual orientation | N | OB | OS | sibs |
| --- | --- | --- | --- | --- | --- | --- |
| Blanchard and Zucker, 1994 | USA. White volunteers from earlier studies<sup>a</sup> | Homosexual | 569 | 286 | 256 | 1104 |
|  |  | Heterosexual | 281 | 123 | 100 | 528 |
| Blanchard and Bogaert, 1996a | Canada : white volunteers<sup>b</sup> | Homosexual | 302 | 213 | 182 | 735 |
|  |  | Heterosexual | 434 | 209 | 206 | 977 |
| Blanchard and Bogaert, 1996b | USA : white individuals from Kinsey database<sup>a</sup> | Homosexual | 799 | 556 | 470 | 1814 |
|  |  | Heterosexual | 3807 | 2223 | 2052 | 8667 |
| Blanchard et al., 1998 | USA, UK : volunteers, from earlier studies<sup>a</sup> | Homosexual | 385 | 205 | 164 | 728 |
|  |  | Heterosexual | 225 | 73 | 96 | 357 |
| Bogaert, 1998 | USA : non-white individuals from Kinsey database. | Homosexual | 229 | 237 | 182 | 810 |
|  |  | Heterosexual | 594 | 630 | 534 | 2370 |
| Ellis and Blanchard, 2001 | USA, Canada : volunteers<sup>a</sup> | Homosexual | 175 | 117 | 85 | 374 |
|  |  | Heterosexual | 971 | 494 | 482 | 1892 |
| Bogaert, 2005 | UK, USA: individuals from prob. sample of households from earlier studies | Homosexual | 79 | 72 | 50 | 204 |
|  |  | Heterosexual | 2721 | 1870 | 1758 | 7249 |
| Blanchard et al., 2006 | Canada : volunteers from previous studies (sample « Bogaert (Other) »)<sup>a</sup> | Homosexual | 267 | 219 | 174 | 655 |
|  |  | Heterosexual | 148 | 75 | 67 | 242 |
| Frisch and Hviid, 2006 | Denmark : national cohort<sup>d</sup> | Homosexual | 1890 | 699 | 594 | 2704 |
|  |  | Heterosexual | 429181 | 147704 | 117529 | 651577 |
| Blanchard and Lippa, 2007 | UK & other : BBC internet survey | Homosexual | 8279 | 4387 | 4235 | 15326 |
|  |  | Heterosexual | 79519 | 35580 | 35368 | 143134 |
| Vasey and VanderLaan, 2007 | Samoa : volunteers | Homosexual | 83 | 188 | 173 | 533 |
|  |  | Heterosexual | 114 | 140 | 142 | 497 |
| Bogaert, 2010 | UK : probability sample from previous study | Homosexual | 132 | 90 | 88 | 331 |
|  |  | Heterosexual | 5472 | 3174 | 3119 | 12148 |
| Schwartz et al., 2010 | USA & Canada : volunteers | Homosexua | 677 | 539 | 445 | 1891 |
|  |  | Heterosexual | 873 | 486 | 446 | 1892 |
| VanderLaan and Vasey, 2011 | Samoa : volunteers | Homosexual | 133 | 255 | 226 | 747 |
|  |  | Heterosexual | 208 | 179 | 212 | 903 |
| Kishida and Rahman, 2015 | UK : volunteers | Homosexual | 905 | 570 | 534 | 1891 |
|  |  | Heterosexual | 999 | 559 | 529 | 2117 |
| Currin et al., 2015 | USA : internet volunteers<sup>a</sup> | Homosexual | 118 | 61 | 57 | 261 |
|  |  | Heterosexual | 500 | 285 | 245 | 1080 |
| Skorska and Bogaert, 2017 | USA : national representative cohort of adolescents, sampled when adult<sup>d</sup> | Homosexual | 225 | 68 | 36 | 289 |
|  |  | Heterosexual | 6562 | 1722 | 1480 | 8675 |
| Xu and Zheng, 2017 | China : internet volunteers | Homosexual | 481 | 118 | 226 | 484 |
|  |  | Heterosexual | 392 | 108 | 164 | 438 |
| Swift-Gallant et al., 2018 | Canada, USA, UK, Australia, New-Zealand : volunteers | Homosexual | 243 | 141 | 122 | 467 |
|  |  | Heterosexual | 91 | 50 | 39 | 191 |
| Nila et al., 2019 | Indonesia : volunteers | Homosexual | 113 | 120 | 140 | 432 |
|  |  | Heterosexual | 62 | 46 | 71 | 210 |
| Apostolou, 2020 | Greece: internet volunteers<sup>b</sup> | Homosexual | 221 | 107 | 87 | 336 |
|  |  | Heterosexual | 593 | 206 | 182 | 850 |
| Gómez Jiménez et al., 2020 | Mexico : volunteers | Homosexual | 244 | 284 | 278 | 855 |
|  |  | Heterosexual | 194 | 145 | 135 | 527 |
| Ablaza et al., 2022 | Netherlands : marriages, population registers<sup>b</sup> | Homosexual | 26542 | 18890 | 16145 | 59984 |
|  |  | Heterosexual | 4607785 | 2795390 | 2687875 | 10874373 |
| This study | France : volunteers | Homosexual | 271 | 128 | 116 | 425 |
|  |  | Heterosexual | 241 | 89 | 100 | 405 |

a. Data available in Blanchard (2018c). b. Data provided upon request. c. Data reconstructed taking into account missing values. d. Data available in Blanchard (2018b).

Two other individual data sets were also considered: the Greek sample described by Apostolou (2020b), and the Indonesian sample used in Nila et al. (2019), and fully described in Nila et al. (2018). For the Greek data, only three categories of men were retained: “exclusively heterosexual”, “bisexual”, and “homosexual” (the category “heterosexual with same-sex attractions” was not considered), and the bisexual category was pooled with the homosexual category (removing bisexual individuals did not change qualitatively the results). Age information was missing for 20 individuals (or 2.5 %), and was replaced by the mean age of the other individuals. For the Indonesian data, the number of maternal aunts and the corresponding number of cousins were also considered. Three individuals were removed due to incomplete data, resulting in a total sample size of *N* = 113. Bisexuals (N = 34) and Waria (a third gender of androphilic males, N = 11) were pooled with the homosexual category (see Nila et al. 2018 for details). For both datasets, individual dates of birth were computed as the year of sampling (June 2018 for the Greek sample) minus the age.

### Statistical analysis of individual family data

#### Generalized Linear models

To assess the presence of AE while controlling for FBOE in the individual family data from France, Greece and Indonesia, two models were considered. First, for the French and Indonesian datasets, a model with the number of cousins from the maternal aunts as the response variable (Model1) tested whether males of different sexual orientation have more or fewer cousins by maternal aunts. Second, a model with the number of siblings as the response variable (Model2) tested whether males of different sexual orientation had more or fewer siblings. For both models, the variable of interest was the sexual orientation of sampled men (non-ordinal qualitative variable), and the control variable was the male birth rank (qualitative variable) of sampled men. Considering male birth rank as a quantitative variable did not qualitatively change the results. The number of maternal aunts was also a control variable for Model1. As the number of cousins or siblings could be influenced by the age of the sampled men, this age (centred and scaled) was added as a control variable for both models. Generalized linear regression was performed, using a Poisson error structure. When the overdispersion parameter (ĉ = residual deviance/residual degrees of freedom) was between 1 and 2, a quasiPoisson error structure was used instead. When ĉ > 2, a Gaussian error structure was used. The significance of each independent variable (explanatory and control variables) was calculated by removing it and comparing the resulting variation in deviance using the *χ²* test (for Poisson or quasiPoisson error structure) or *F* test (for Gaussian error structure), as done by the function *Anova* from the *car* R package.

#### Bayesian modelling

In order to estimate the relative contribution of both FBOE and AE on number of siblings, we performed a Bayesian analysis using a hierarchical model, implemented in the NIMBLE R package (de Valpine et al., 2017). The birth order effect, i.e. the probability *p* of displaying a homosexual preference according to the number of older siblings was modelled as *p*=*f* (*p*_0_ *, X*) where X is the number of older brothers (for modelling an older brother effect), or the number of older sisters (for modelling an older sister effect), and *p_0_* is the probability of a firstborn displaying a homosexual preference (thus p0 is also the probability of sampling a homosexual among the first born individuals from the population sample considered). Various forms of the function *f* were considered, notably logistic, geometric, linear, and polynomial (Table S1). These functions describing the FBOE effect were compared by evaluating model fit to the sibling data using WAIC (Watanabe-Akaike Information Criterion, a generalized version of AIC onto singular statistics models, see Gelman et al. (2014) and Watanabe (2013)): the mean of ten independent chains was used, each with a length of 50,000 samples and a burn-in phase of 20,000. To avoid the effects of small sample sizes for the number of older brothers or sisters in categories poorly sampled, we restricted the data, for the WAIC comparison, to categories including at least 10 individuals. Fertility was assumed to follow a Poisson distribution with rate parameter λ. Three types of heterogeneity in *λ* were simultaneously considered. First, a temporal variation of λ over the last decades: log(*λ)* was modelled as a linear function of the year of birth (*yob,* continuous variable, centered and scaled) of the sampled individuals, *λ*=*e^c^*^1+^*^c^*^2.^*^yob^*. Second, the possible presence of a subgroup displaying a higher fertility was modelled with parameter *h* (with *h*≥0). For each individual, the probability that modified fertility rate parameter *λ(1+h)* applies followed a Bernoulli distribution with parameter *φ*. Third, the possible presence of a subgroup displaying a higher fertility and simultaneously a larger value of *p_0_*. This antagonist effect was modelled as an increase in *λ* by a factor 1+ *β*, conjointly with an increase in *p_0_* by a factor (1+ *α*) (with *α* and *β*≥0). For each individual, the probability that the AE applies followed a Bernoulli of parameter *φ*_ae_. We implemented the model in a Bayesian framework by assigning uninformative prior distributions for all model parameters, and fitting the model using Markov chain Monte Carlo (MCMC) in the nimble R package. The birth order effect and the antagonist effect were simultaneously modelled in Nimble (de Valpine et al., 2017). A chain with a length of 100,000 samples and a burn-in phase of 10,000 was used to compute the posterior distributions of the parameter estimates. Support for each effect (AE or FBOE), in the presence of the other one, was computed using Reversible-Jump Markov chain Monte Carlo (or RJMCMC, Green, 1995). RJMCMC is an extension to standard MCMC methodology that allows simulation of the posterior distribution on spaces of varying dimensions. Toggle samplers controlled the inclusion or exclusion of each effect according to RJMCMC transition probabilities. Two indicator variables controlled the presence or absence of the AE parameters (*α* and *β)*, and one indicator variable dictated presence or absence of the FBOE or SBOE parameter (*a_1_*). RJMCMC was run at least 200,000 iterations, and the mean of the posterior distribution of the binary inclusion variables for each effect were used as an estimate of the support of the effect considered. When posterior samples of the binary inclusion variables appeared to have not converged to a stationary distribution, thus decreasing confidence in the posterior results, longer chains were applied (i.e. 2×10^6^ iterations). Analyses were run in R 3.6.3 (R Core Team, 2020) using version 0.9.0 of the nimble package (de Valpine et al., 2020).

## RESULTS

### Methodological developments

#### 1) Relationship between mean birth rank and mean fertility

Let us consider a population of *N* families with discrete generations, each family having a number of children drawn from a Poisson distribution of parameter *λ*. Among the *λN* expected children, *N* (1−*e*^−^ *^λ^*) are firstborn and *N* (1−*e*^−^ *^λ^*−*λe*^−^*^λ^*) are second born. The probability of sampling an individual with birth order *j ≥* 1 is (Appendix 1):

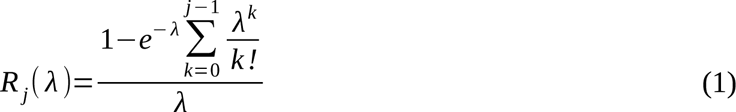

The expected value 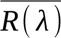 of this sampling distribution is (Appendix 1, Supplementary materials):

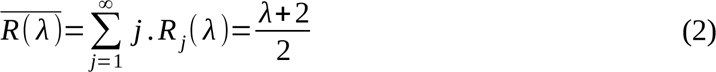

Simulation was used to verify equation (2) for *λ* values from 0.5 to 6 (Fig. S1).

#### 2) Estimating fecundity from sibling data

There are two known biases when fecundity is estimated from sibling data (Keyfitz and Caswell, 2005). First, mothers with no children cannot be sampled with this method, thus inflating fertility estimates. A zero-truncated sampling distribution is thus required. Second, the probability of sampling a member from a sibship class of any given size in the general population is proportionate to the number of siblings (review in Berglin, 1980). A correction for this second bias was proposed in 1914, but “*it is usual to find that authors are unacquainted with the trap*” (Berglin, 1980). Both biases lead to an overestimate of fecundity, this overestimate being predominant for low fecundity values for the first bias (because the probability of sampling the zero-class becomes relatively high), and for high fecundity for the second bias (because the variance in sibling size increases). An unbiased estimate of population fecundity from sibling data, taking into account both sources of bias, and considering that fecundity follows a Poisson distribution, is given by the mean number of siblings of the sampled individual (mean number of brothers and sisters, not including the sampled individuals), see Appendix 2, Supplementary materials. Simulation was used to check the various corrections proposed, for *λ* values from 0.5 to 6 (Fig. S2). For aggregated data (population samples), the mean fecundity is given by the total sibship size divided by the number of sampled individuals (as 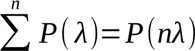), where *P*(*λ*) is a Poisson distribution of parameter *λ*).

#### 3) Sampling distribution of number of older brothers or older sisters

When men are randomly sampled in the absence of FBOE (and without taking into account sexual orientation) the sampling distribution of the number of their older brothers (*ob*) is given by Prob(*ob* = *i*) = *R_i+1_*(*λ* /2), where *R* is from Eq. (1), and *λ* /2 is the fertility considering only male offspring (i.e. half the overall fertility, assuming a balanced sex-ratio). The probability distribution of older sisters and older brothers should be the same, unless some male birth rank categories are over or under represented during the sampling of homosexual men (as would be the case when FBOE is operating). Thus, in the absence of FBOE, the probability distribution of older sisters Prob(*os* = *i*) could be also calculated considering that men of various male birth orders are sampled, giving (see Appendix 1b for derivation):

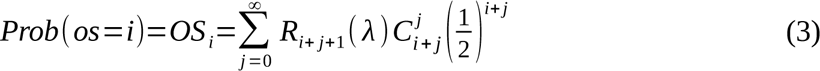

#### 4) Simulating SBOE and FBOE

In the absence of FBOE, the sampling distribution of the number of older sisters is given by Eq. 3. An FBOE will lead to under-representation of low birth rank categories (e.g. first born) and over-representation of high birth rank categories in samples of homosexual men, changing the R_i_(*λ* /2) and R_i_(*λ*) values, thus affecting the sampling distribution of older sisters of Eq. 3. The sampling distribution of the number of older sisters in presence of an FBOE is not easily tractable analytically, thus simulation was used to assess if an FBOE generates an apparent SBOE. Five thousand families were generated, with mean fecundity 4 and 1:1 sex-ratio. An FBOE was modelled by considering that the probability of being homosexual for *i* > 0 older brothers is increased by a constant *a* proportional to *i* (function f5, Table S1), or otherwise increased by a constant *a1* (function f7, Table S1), with *a* = *a_1_* = 0.2. From a random sample of 500 heterosexual and 500 homosexual men, the proportion of homosexual men was computed for each older brother or older sister category. The mean of 1000 replicates of this process was computed, with FBOE modelled using functions f5 or f7, or without FBOE as a control. A substantial older sister effect appears when randomly sampling hetero- and homosexual men in the presence of FBOE only (Figure 1, A and B). In absence of an FBOE no SBOE is observed from the same sampling process (Figure 1, C).

**Figure 1.**
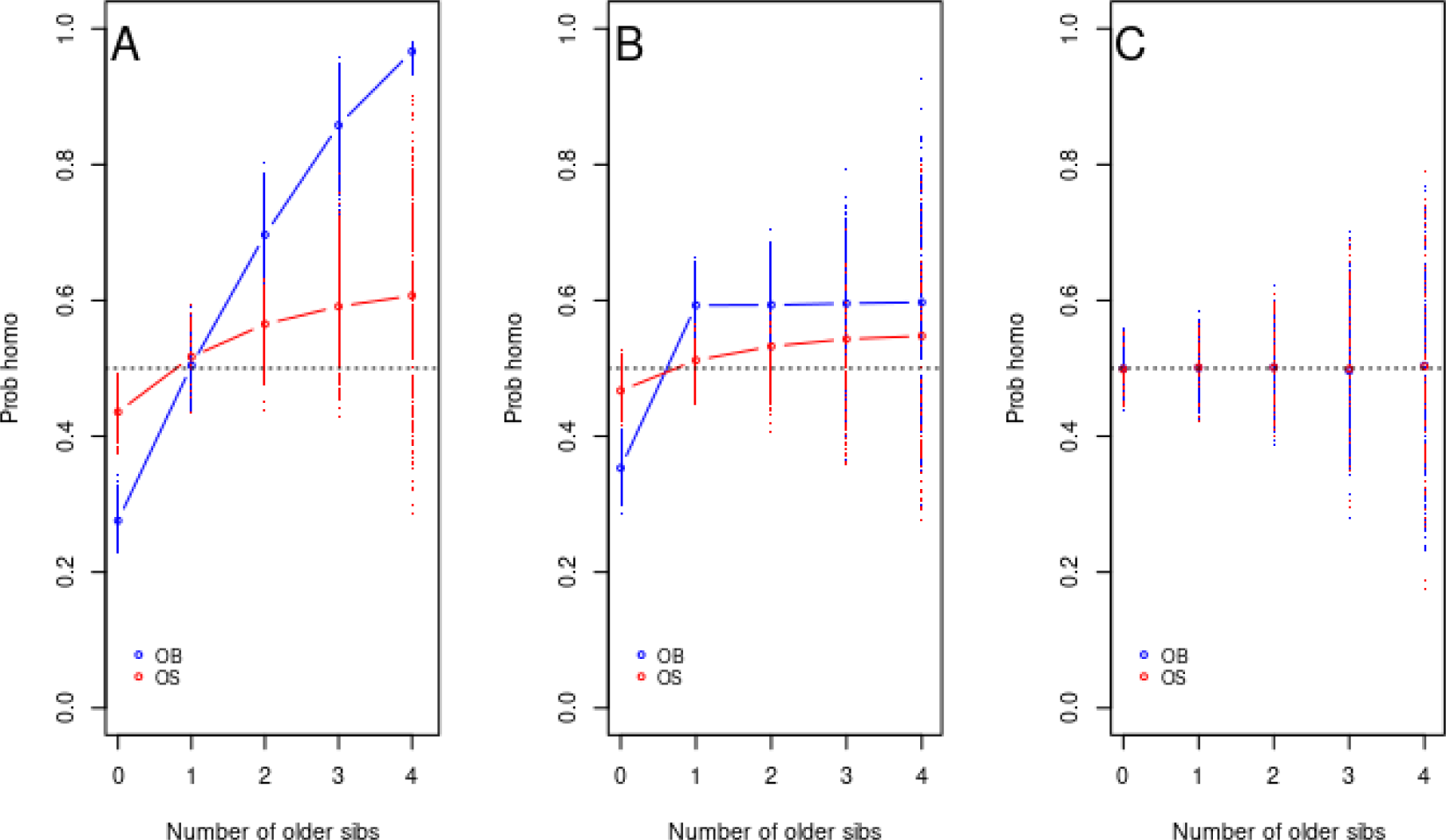
Apparent SBOE when sampling men from a population with only an FBOE. A) FBOE generated with function f5 (cf Table S1). B) FBOE generated with function f7. C) Control (no FBOE). For each of the 1000 replicates, 500 heterosexual and 500 homosexual men were randomly sampled, and the proportion of homosexual men was computed for each older brother (blue) or older sister (red) category and is represented as a dot. The mean for these replicates, for each category of older siblings, is depicted as an empty circle with the corresponding colour. The dotted black line represents the expected curve when sexual orientation is independent of male birth rank.

### Empirical data analysis

#### Aggregated family data

A total of 23 pairs of samples of aggregated data was retained from the published literature, thus representing, with the French dataset, a total of 43,362 homosexuals and 5,141,967 heterosexuals (Table 1). Mean fertility of mothers ranged from 1.0 to 6.4 for homosexual samples, and from 1.1 to 4.4 for heterosexual samples. The mean birth rank of men with respect to only their brothers (computed as *OB/N +1*, see above) was between 1.2 and 3.3 for homosexual samples, and between 1.3 and 2.2 for heterosexual samples. For heterosexuals, the relationship between mean number of male offspring in the sibship (i.e. mean number of sons) and mean male birth rank did not deviate from the theoretical prediction (Figure 2): slope = 0.497 (SE = 0.04), not significantly different from the expected value of 0.5 (*F_1,21_* = 0.007, *P* = 0.93), and intercept = 0.998 (SE =0.04), not significantly different from the expected value of 1 (*t*(22) = −0.04, *P* = 0.97). For homosexuals, this relationship displayed a slope of 0.72 (SE = 0.02), which is significantly higher than 0.5 (*F_1,22_* = 85.5, *P* < 10^-8^).

**Figure 2.**
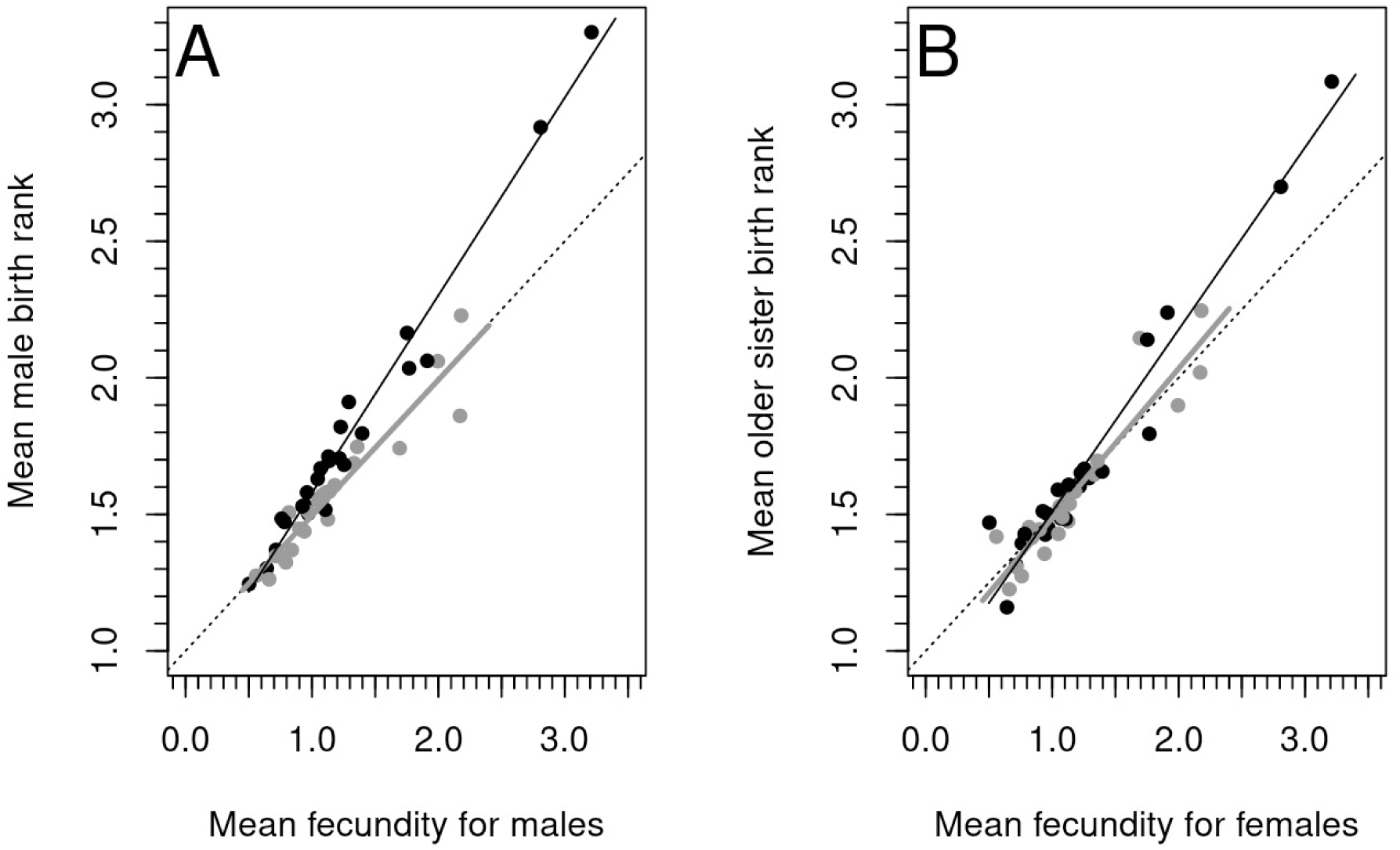
Mean birth rank in relation to fecundity. A. Birth rank among males. B. Birth rank among older sisters. The mean fecundity for only males or only females is computed as half the overall fecundity. Each sample is represented as a dark dot for homosexuals, and as a grey dot for heterosexuals. The solid lines are regression lines for homosexual samples (black), and heterosexual samples (grey). The dotted line represents the theoretical expectation between mean birth rank and mean fecundity (i.e. Equation 2).

The mean birth rank of men with respect to only their sisters (computed as *OS/N +1*, see above) was between 1.2 and 3.1 for homosexual samples, and between 1.2 and 2.2 for heterosexual samples. For heterosexuals, the relationship between mean number of female offspring in the sibship (i.e. expected number of sisters) and mean birth rank of men with respect to only their sisters was not different from the theoretical prediction: slope = 0.55 (SE = 0.05), not significantly different from the expected value of 0.5 (*F_1,22_* = 1.17, *P* = 0.30), the intercept = 0.94 (SE = 0.05) was not significantly different from the expected value of 1 (*t*(22) = −1.1, *P* = 0.27). For homosexuals, this relationship displayed a slope of 0.67 (SE = 0.03), significantly higher than 0.5 (*F_1,22_* = 23.8, *P* < 10^-4^).

Data simulation was used to decipher which phenomenon could generate such a higher slope for the relationship between mean number of sons and mean birth rank of men among brothers for homosexuals. When an AE alone was modelled, the resulting slope for homosexuals or heterosexuals were not higher (*P* > 0.5) that the theoretical value of 0.5 (Table S2). When an FBOE alone was modelled, a significantly (*P* < 0.001) higher slope was observed when the probability of being homosexual increased linearly with the number of older brothers (function f5, Table S1) or, for a threshold function, when the effect of having at least one older brother increased with the mean fertility (function f7’) (Table S2, Figure S3). When AE and FBOE were simultaneously modelled, results were globally similar to those with an FBOE only. Thus, a slope larger than the expected value of 0.5 suggests the presence of an FBOE, and is not informative for the presence of AE. Using the aggregated family data, maximum likelihood estimate of the parameters were â = 0.240, SE = 0.23 (function f5), and μ = 0.152, SE = 0.14 (function f7’).

#### Individual family data

For the three datasets, the number of older brothers, older sisters, and siblings are given in Table 1, and descriptive statistics are given in Table 2. All datasets displayed a higher number of older brothers and older sisters for homosexual men (for the Indonesian sample, see Nila et al., 2019), and this difference was significant (Wilcoxon Mann-Whitney, *P* < 0.05) except for the French dataset (Older brother: *P* = 0.10; older sisters: *P* = 0.74). For the Greek and Indonesian samples, maternal fertility was significantly higher (Wilcoxon Mann-Whitney, *P* < 0.05) for homosexuals than heterosexuals. For the French sample, heterosexuals displayed a non-significantly (*P* = 0.49) higher maternal fertility. When controlled for birth rank, the fertility of mothers was not different in homosexuals compared to heterosexuals (Indonesia: *P* = 0.60; France: *P* = 0.32; Greece = 0.93, Table S3). The same result was found for the fertility of aunts: the number of cousins, controlled for the number of aunts, was not different between homosexuals and heterosexuals, for the same birth order (Indonesia: *P* = 0.82; France: *P* = 0.08, Table S4).

**Table 2.** Descriptive statistics for the three population samples of individual family data. The Greek and Indonesia samples are from Apostolou (2020) and Nila et al. (2019), respectively. Numbers of OB, OS, and siblings are in Table 1.

|  | Indonesia |  |  | Greece |  |  | France |  |  |
| --- | --- | --- | --- | --- | --- | --- | --- | --- | --- |
|  | Homo | Hetero | All | Homo | Hetero | All | Homo | Hetero | All |
| N | 113 | 62 | 175 | 221 | 593 | 814 | 271 | 241 | 512 |
| Mean age (SD) | 31.6<br>(9.4) | 30.0<br>(12.6) | 31.0<br>(10.6) | 29.8<br>(8.5) | 35.1<br>(11.9) | 33.7<br>(11.3) | 32.0<br>(10.0) | 35.8<br>(13.4) | 33.8<br>(11.8) |
| Mean year of birth | 1983.4 | 1985.0 | 1984.0 | 1988.7 | 1983.4 | 1984.8 | 1978.5 | 1973.1 | 1976.0 |
| Mean OB | 1.06 | 0.74 | 0.95 | 0.48 | 0.35 | 0.38 | 0.47 | 0.37 | 0.44 |
| Mean OS | 1.24 | 1.15 | 1.21 | 0.39 | 0.31 | 0.33 | 0.43 | 0.41 | 0.42 |
| Mean sibs | 3.82 | 3.39 | 3.67 | 1.52 | 1.43 | 1.46 | 1.56 | 1.68 | 1.62 |
| Aunts | 221 | 124 | 345 | - | - | - | 394 | 288 | 682 |
| Cousins | 535 | 316 | 851 | - | - | - | 794 | 497 | 1291 |

Bayesian inference was used to test whether an AE, or an FBOE (or an SBOE), or both, could be inferred from these individual family data. For each dataset, the various functions (Table S1) describing the variation of probability of a homosexual orientation according to the number of older siblings (brother or sisters) were compared using WAIC. For an FBOE, the function providing the minimum WAIC (or WAIC_min_,) were f7 (for France and Indonesia) and f5 (for Greece). For all datasets, these two functions provided a WAIC value lower than WAIC_min_+2 (Table S5). For an SBOE, the minimum WAIC value resulted from function f7 (for Indonesia and Greece) and function f2 (for France). Only two functions (f4 and f7) provided a WAIC lower than WAIC_min_+2 for all datasets (Table S5). Thus function f7 was chosen to describe the older sibling effect for further modelling. This function fits two parameters, the probability, in the dataset, of sampling a homosexual with no older siblings (*p_0_*), and the increase (*a_1_*) of this probability for one or more older sibling, thus describing a constant sibling effect starting with the first older sibling (Figure 3).

**Figure 3.**
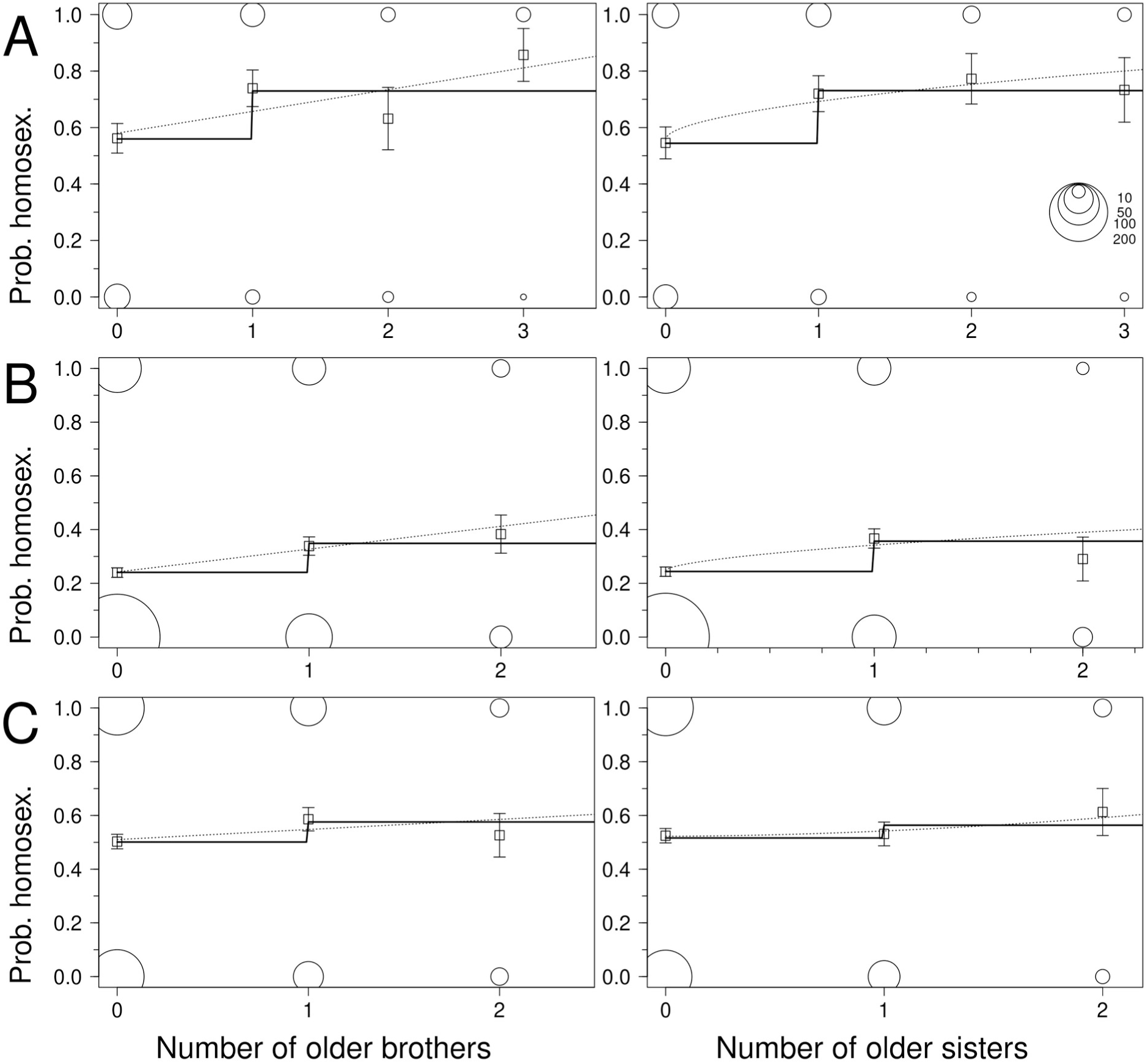
Modelling the older sibling effect. Data for the three datasets are presented (A: Indonesia; B: Greece; C: France), for each type of older siblings (left: older brothers; right: older sisters). The solid lines correspond to function 7. The dotted lines correspond to function 5 (left panels) or function 4 (right panels). The number of observations for each older older sibling category (at 0 on the y axis for heterosexuals, and at 1 on the y axis for homosexuals) is represented as a circle with an area proportional to sample size, according to the reference legend in the top-right panel. The frequency of homosexuals is depicted as a square, with the corresponding +/− SE range. Data with elevated number of older siblings (A: more than 3; B-C: more than 2) are not represented.

For each dataset, sibling data were fitted for an AE simultaneously with an FBOE or with an SBOE. Fertility was fitted to take into account two possible sources of heterogeneity (variation of fertility with year of birth, and a subgroup of individuals with a different fertility), in addition to a possible AE. Means of the posterior distribution of the parameters are presented Table 3, and are used to estimate λ and its variation across detected groups, and the importance of AE. In the Indonesian sample, the mean fertility was *λ*=*e^c^*^1^ =2.25 in 1983.2 (mean year of birth), and decreased to 1.66 ten years later (or 0.94 standard deviations later). A group representing 19% of individuals (Bernoulli parameter φ = 0.19) displayed only a higher fertility of *λ*(*1+h*) = 7.72, and another group representing 27.6 % of individuals (Bernoulli parameter φ_ae_= 0.27) displayed a higher fertility *λ*(*1+β*) = 5.54 and at the same time an increased probability of being homosexual among the first born in the dataset (*p_o_*(*1+α*) = 0.84). In the Greek sample, two groups of individuals were identified, one representing 54 % of individuals (Bernoulli parameter φ =0.54) with a mean fertility of *λ*=*e^c^*^1^ =1.02, and another (1-φ = 46%) with a mean fertility of *λ*(*1+h*) = 2.96. No substantial temporal variation was detected. The AE concerned 25% of individuals (Bernoulli parameter φ_ae_= 0.25), whose probability of being homosexual among the first born in the dataset was *p_o_*(*1+α*) = 0.32, and displaying a fertility of *λ*(*1+β*) = 3.84. In the French sample, two groups of individuals were identified, one (Bernoulli parameter φ =0.55) with a mean fertility of *λ*=*e^c^*^1^ =0.76, and another (1-φ ∼ 0.45) with a mean fertility of *λ*(*1+h*) = 3.28. A decrease of the mean fertility with time was apparent, from (computed for the fraction *λ*(*1+*h)) = 3.28 in 1976.0 (mean year of birth), to 3.0 ten years (or 0.76 standard deviations) later. The AE concerned 49.2% (Bernoulli parameter φ_ae_= 0.49) of individuals, whose probability of being homosexual among the first born in the dataset was *p_o_*(*1+α*) = 0.73, and displaying a fertility of *λ*(*1+β*) = 1.88 and 8.13, for the two groups above, respectively.

**Table 3.** Parameters estimates when AE and FBOE are simultaneously considered. Mean and standard deviation (SE) of the parameter posterior distribution. The proportion of individuals concerned (*φ* and *φ*_ae_) is indicated for parameter values dependent on a latent variable. The variation in time is described by the intercept (*c_1_*) and a slope (*c_2_*). See text for interpretation of the parameters.

|  |  | Indonesia |  | Greece |  | France |  |
| --- | --- | --- | --- | --- | --- | --- | --- |
|  |  | Mean (SE) |  | Mean (SE) |  | Mean (SE) |  |
| Fertility |  |  |  |  |  |  |  |
| Variation in time : |  |  |  |  |  |  |  |
|  | c1 | 0.810 | (0.506) | -0.105 | (0.588) | -0.271 | (0.801) |
|  | c2 | -0.321 | (0.044) | 0.015 | (0.032) | -0.117 | (0.034) |
| Hyperfertility : |  |  |  |  |  |  |  |
|  | h | 2.435 | (1.553) | 2.286 | (2.285) | 3.297 | (2.212) |
| | $\varphi$ | 0.190 | (0.294) | 0.460 | 0.426 | 0.446 | (0.404) |
| AE : |  |  |  |  |  |  |  |
| | $\alpha$ | 0.685 | (0.584) | 0.492 | (0.506) | 0.853 | (0.589) |
| | $\beta$ | 1.464 | (1.350) | 3.193 | (1.931) | 1.483 | (2.272) |
| | $\varphi_{ae}$ | 0.276 | (0.311) | 0.249 | (0.391) | 0.492 | (0.411) |
| OBE |  |  |  |  |  |  |  |
|  | p0 | 0.498 | (0.098) | 0.215 | (0.048) | 0.392 | (0.110) |
|  | a1 | 0.155 | (0.311) | 0.104 | (0.035) | 0.082 | (0.042) |

Support values for FBOE or SBOE (in presence of AE), and for AE (in presence of FBOE or SBOE) were evaluated using RJMCMC. In the presence of an older sibling effect, either FBOE or SBOE, there was little support for an antagonist effect (Figure 4). The maximum support values were ∼20% for the Indonesian dataset. In the presence of AE, there was a large support for an FBOE or SBOE, in the Indonesian and Greek datasets (all supports > 50%). For the French dataset, support was limited (∼30%), or non-existent (<1%), for FBOE, and SBOE, respectively (Figure 4).

**Figure 4.**
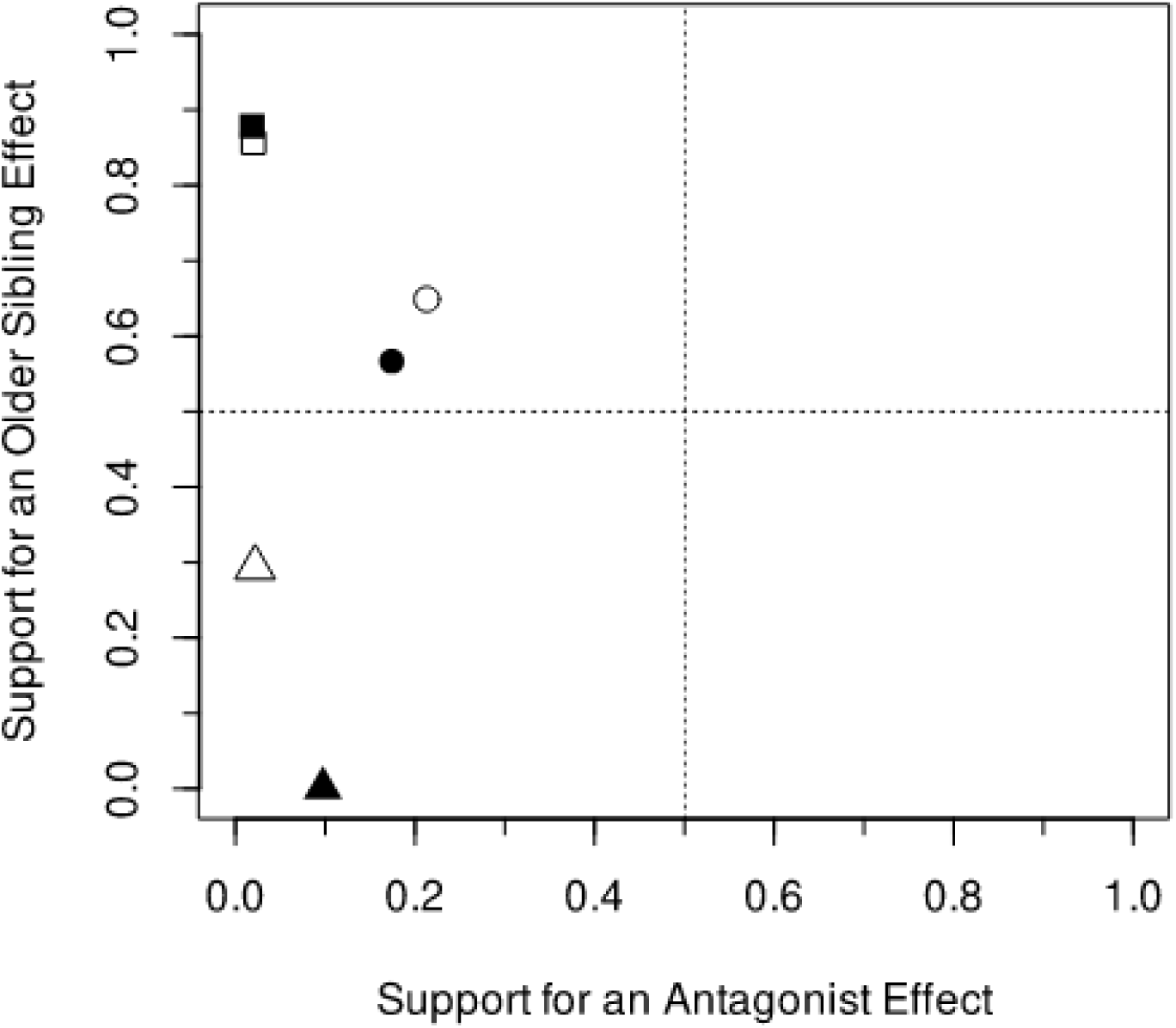
Support for an antagonist effect (AE) or an older sibling effect, conjointly modelled. Coordinates of each point represents the support for AE in presence of an older sibling effect (x-axis), and the support for an older sibling effect in presence of an AE (y-axis). The older sibling effect is either an older brother effect (FBOE, full symbol), or an older sister effect (SBOE, open symbol). Point shapes vary according to datasets (circle, square, and triangle for Indonesia, Greece, and France, respectively).

## DISCUSSION

Research on the biological determinants of male homosexual preference have long realized that the older brother effect (FBOE) and the antagonist effect (AE) can both generate family data where male homosexual men have more siblings, more relatives and more older siblings than heterosexual men. Here, we developed several approaches to disentangle these two mechanisms from empirical population samples or family samples. By analysing three types of datasets with statistical tools correcting for known sampling biases, we were able for the first time to separately test the actions of the birth rank and antagonist genetic factors on fertility and sexual orientation. We found unambiguous support for the FBOE in aggregated population data from 24 independent samples, as well as in two individual datasets out of three. We showed that an apparent SBOE can be generated by sampling bias in presence of an FBOE, and conclude that the SBOE reported in some previous studies is probably artefactual. We found no support for the AE in individual datasets including the extended maternal family. Levels of statistical support for FBOE and/or AE, in the various datasets, are shown in Table 4.

**Table 4.** Results of the various tests for detecting the presence of FBOE or AE in the different datasets considered. A dash indicates that the hypothesis could not be tested.

| Type of data | Hypothesis tested |  |
| --- | --- | --- |
|  | OBE | AE |
| Aggregated data | Yes | - |
| Individual data |  |  |
| Indonesia : |  |  |
| Mother fertility | - | No |
| Aunts fertility | - | No |
| siblings | Yes | No |
| Greece : |  |  |
| Mother fertility | - | No |
| siblings | Yes | No |
| France : |  |  |
| Mother fertility | - | No |
| Aunts fertility | - | No |
| siblings | No | No |

### Sampling biases in presence of an older brother effect generate an artefactual older sister effect

*“The history of birth order studies is not a happy one*”. The warnings mentioned by Price and Hare (1969), and regularly recalled since then (e.g. Berglin, 1980; Keyfitz and Caswell, 2005), correspond to several sampling biases, identified long since, but not always taken into account, while the generation of various statistics to test various hypotheses has added to the confusion. For example, indices proposed to study SBOE have not considered sampling biases generated when FBOE is present in the population. The explanation is simple: individuals with more older brothers are more often sampled from larger sibship size (i.e. from a mother with a higher fertility), thus with also more older sisters, considering an even sex-ratio. The sampling distribution of the number of older brothers or sisters in a sample of men (Eq. 1 and 3) is a first step towards developing adequate statistics, although an explicit form of FBOE should be introduced to define the sampling distribution of the number of older sisters in presence of FBOE. The correlation between the number of older brothers and the number of older sisters has been widely acknowledged previously (e.g. Blanchard and VanderLaan, 2015), but not sufficiently considered. This sampling bias does not rule out the action of a genuine SBOE in population data, but any claim for an SBOE, or for any additional sibling effect, should first control for the sampling bias generated by FBOE. Thus, the report, in a recent meta-analysis, of a widespread SBOE in addition to the FBOE (Blanchard et al., 2021), should be treated with caution. Blanchard and Lippa (2021) present some evidence for an SBOE using the large dataset of Blanchard and Lippa (2007) by selecting individuals with only one sib (thus controlling for fertility) and without older brothers (thus controlling of the FBOE): one older sister, relatively to a younger sib, significantly increases the probability of homosexuality (*P* = 0.02), although a re-analysis using a more rigorous test (one older sister vs a younger sister) would have given *P* = 0.04 (details in Appendix 3, Supplementary Materials). This provides significant, although weak, support for a SBOE. Ablaza et al. (2022) also provide some support for a genuine SBOE, although their new regression method, using several highly correlated variables, requires a formal validation. We thus agree that empirical data reveal a genuine SBOE, although a thorough validation is required.

### The older brother effect is well supported but its effects depend on mean population fertility

The generality of the FBOE has generated much recent debate fuelled by analyses of aggregated data (Blanchard, 2018c; Blanchard et al., 2021; Vilsmeier et al., 2021). Our re-analysis of all available and relevant aggregated data strongly supports the generality of the FBOE, and rejects that it is an artefact from AEs. We provide several improvements to clarify the phenomenon. First, we filtered out samples not corresponding to adult typical homosexuality, such as pedophiles, or corresponding to non-representative populations, such as sex offenders, transexuals, psychoanalytic or hospital patients (Zietsch, 2018). Thus, our results can be safely associated with standard homosexual men, and it would be interesting to test whether FBOE applies more generally. Second, we derived the relationship between mean birth rank and mean fertility in population samples, and showed in simulated data that the slope of this relationship was changed for homosexual men by an FBOE but not by an AE: an FBOE generates different slopes for homosexual and heterosexual samples while an AE moves the homosexual sample towards higher fertility but with the same slope as heterosexual men (Table S2, Fig. S3). We then showed that population data from heterosexual men did not deviate from the expected relationship between birth rank and mean fertility while data from homosexual men showed a steeper slope (Fig 2A), which can thus be safely attributed to an FBOE rather than to any confounding fertility effect such as the AE. This also suggests that the expression of the FBOE is fertility-dependent, as the probability of being homosexual increases with fertility, and two processes may generate this effect. First, the probability of being homosexual increases with male birth order, and mean male birth order increases with fertility (function f5 in Table S2). In this case FBOE is not stronger when fertility increases but this trait is more often expressed due to an increased number of larger families. Alternatively, the probability of being homosexual is constant for one or more older brothers, but this constant increases with mean fertility (function f7’ in Table S2). In this case, FBOE is stronger when mean fertility increases. At present we cannot distinguish between these two possibilities.

The fact that the expression of FBOE is fertility-dependent has a consequence for the study of FBOE. When indices are used to control for fertility, such as those proposed by Slater (1962), Berglin (1980), Blanchard (2014; 2018b; 2018c), or Vilsmeier et al. (2021), the implicit assumption is that homosexual and heterosexual samples are compared independent of the mean fertility level, and the variation of those indices as a function of the mean fertility is therefore not evaluated. When the mean fertility is low (e.g. λ ≤ 2, thus mean number of sons ≤ 1), the mean rank of homosexuals is not very different from that of heterosexuals, and the FBOE effect is not apparent (see Fig 2A). This probably explains the reports of an absence of FBOE in single-sample studies from low fertility populations, e. g. from France (this study), and UK (Kishida and Rahman, 2015; Rahman et al., 2008). Thus, in a meta-analysis, including samples from populations with various fertility values, results will probably depend on the number and size of samples from low or high-fertile populations, where FBOE is differently expressed, unless the fertility-dependent expression of FBOE is explicitly considered.

### The shape of the older brother effect remains elusive

At which rate additional older brothers increase the probability of homosexuality is not known. Our data did not permit us to distinguish among the five functions proposed (f3 to f7, Table S1), including a logistic, geometric, and linear functions (Table S5). This low resolution is explained by the paucity of individuals displaying a relatively high number of older brothers (e.g. ob > 3), thus precluding distinction of the various functions. We know of only one previous attempt to infer the shape of the FBOE: Cantor et al. (2002) used the data from Blanchard and Bogaert (1996a) and Blanchard et al. (1998) to fit linear and quadratic functions, and found no significant support for including terms of degree > 1. A linear relationship between number of older brother and probability of being homosexual has been generally assumed since this seminal work. However, such an unbounded linear increasing function is necessarily an approximation useful only for cases with few older brothers. Similarly, the threshold function f7 used here certainly represents another approximation, also poorly applicable to large sibships with many older brothers.

### Proximate and ultimate mechanisms of the older brother effect

The main candidate for a proximate mechanism for the FBOE is a maternal immune response to male-specific antigens (Blanchard and Bogaert, 1996b; Bogaert and Skorska, 2011), for which possible molecular evidence has been recently presented (Bogaert et al., 2018). Exclusive same-sex preference is not found among non-human primates, thus it is not possible to evaluate whether FBOE is restricted to humans. However, based on current knowledge, effect of birth order on sexual orientation is only found in humans, and in no other primate species including close human relatives. This suggests that the FBOE is not a mere constraint of the gestation in primates, and thus the effect of male birth order on sexual orientation requires an evolutionary explanation. Under this hypothesis, the FBOE would be an adaptive plastic manipulation of the phenotype of male offspring by the mother. Nila et al. (2019) have proposed that the FBOE could decrease male sibling competition by later-born males. Such a mechanism could be selected for in patrilocal societies, but probably not in matrilocal ones, where males usually migrate, thus reducing local competition. It should be then interesting to compare FBOE between patri- and matrilocal societies. More generally, the relative contribution of adaptive responses and developmental constraints in shaping the FBOE, and the selective pressures generating the FBOE, remain in urgent need of investigation.

### No support for antagonistic pleiotropy through female fertility

After controlling for the confounding effect of the FBOE on fertility in families of heterosexuals and homosexuals, we found no direct association between higher maternal fertility and male homosexual orientation, i.e. no support for genetic factors increasing fertility of females and increasing at the same time the probability that any given son is homosexual. The larger sibship size displayed by homosexual men is indeed best explained by their high male birth rank (or FBOE). More fertile women are more likely to produce homosexual sons because they are more likely to produce sons with a high birth rank (thus with several older brothers), and not because they have a higher propensity to produce homosexual sons at any given birth rank, compared to lower fertility women. Sampling homosexual men, randomly relatively to their birth rank, will thus result in individuals with a higher number of older brothers due to FBOE, but also with a larger number of siblings. These highly fertile mothers are likely to have sisters also displaying high fertility, due to correlation of fertility among sisters (Anderton et al., 1987; Berent, 1953). We found no difference in mother or aunt fertility between homosexuals and heterosexuals after controlling for male birth rank. In addition, we found no evidence for AE in the analysis of individual datasets, including one displaying a high fertility (Fig. 3).

The antagonist pleiotropy hypothesis proposes that the reproductive cost of homosexual men is at least counter-balanced by a reproductive advantage of relatives, and that both effects are driven by the same genetic factors. If the advantage is greater than the cost, then those genetic factors increase in frequency, and the frequency of homosexual men increases in the population up to the point where the cost becomes too high (if the fitness loss of males is sufficiently large, leading to protected polymorphism, see Gavrilets and Rice, 2006). It was first proposed that maternal female relatives were concerned (e.g. mothers and aunts), and expressed a higher fecundity (Camperio-Ciani et al., 2004b). The higher fecundity of maternal female relatives was subsequently also found in several independent datasets (Iemmola and Camperio-Ciani, 2009; Rahman et al., 2008; Vasey and VanderLaan, 2007) However, these analyses did not control for FBOE, possibly leading to artefactual results due to the sampling biases we have described above. Unfortunately, the original data used by Camperio-Ciani and collaborators to test for the AE are no longer available (Camperio-Ciani pers. comm., February 2020), and could not be reanalysed. Until additional data or additional analyses are presented, we suggest that there is currently no evidence for antagonist pleiotropic factors that compensate reduced reproductive success of homosexual sons with higher fecundity of their female relatives.

Zietsch et al. (2008) were the first to suggest that the a compensatory fitness advantage of genetic factors increasing the occurrence of homosexuality could be expressed by heterosexual relatives of both sexes, in the form of a higher number of sex-partners. The idea is that genes predisposing to homosexual orientation may also increase mating success in heterosexuals. This hypothesis has recently received empirical support from genomic evidence for such pleiotropic genes (Zietsch et al., 2021), where the pleiotropic advantage associated with male or female homosexuality seemed to be restricted to a mating advantage of heterosexual male relatives (Table S2 of Zietsch et al., 2021). Reproductive output is notoriously difficult to measure for males, due to variable mating strategies and extra-pair copulation, although paternity uncertainty is low in some human populations (Larmuseau et al., 2016, 2016; Larmuseau et al., 2019). Further studies are required to confirm the presence of such an intra-sex antagonist effect. Interestingly, the cross-sex genetic correlation for male homosexuals (i.e. between male homosexuals and number of children for female relatives) in Zietsch et al. (2021) was non-significant, consistent with the absence of AE for female fertility that we report here.

### Limits and future directions

This study has several limitations, although none of these call our results into question. First, the aggregated data mainly rely on published studies, thus generating a potential publication bias (excess of publication with significant FBOE). However, the new method of analysis proposed here should not be very sensitive to such publication bias, as it mainly relies on samples displaying different fertilities (in order to estimate the slope of the increase of mean birth rank as a function of mean fertility). A larger proportion of studies from high-fertility population would nevertheless strengthen the results. The specific case of stopping rules, influencing parents in deciding whether to have another child depending of the sex of the previous ones, has not been specifically considered. This phenomenon could affect older siblings’ sex composition and thus the ability to detect FBOE in low fertility populations (Blanchard, 2022). Second, only three individual datasets were analysed, thus restricting the generality of the results concerning the joint analysis of FBOE and AE. Third, a 1:1 male/female sex ratio at birth has been assumed when deriving the sampling distribution of older sisters, whereas the commonly observed sex ratio is approximately 105 boys born per 100 girls. This slightly male biased sex-ratio should be taken into account for more precise derivation. Fourth, bisexual individuals were pooled with homosexuals, as this is commonly done (for a recent example, see Blanchard and Lippa, 2021), even if there is no formal justification. Nevertheless, if bisexuals are closer to heterosexuals than to homosexuals for the traits under study, pooling together bisexuals and homosexuals should make any significant result more conservative, and non-significant results questionable. Here, some non-significant results were found for the individual dataset (see Table 4), but removing the bisexual individuals did not changed them qualitatively (details not shown). Finally, modelling assumed that maternal fertility in human populations follows a Poisson distribution, which is not always the case (e.g. Austerlitz and Heyer, 1998; Hruschka and Burger, 2016). Even if several sources of additional fertility variability have been explicitly incorporated, alternative probability distributions, such as the negative binomial, should be also considered.

## ACKNOWLEDGEMENTS

We thank Maxime Derex for collecting data for the French dataset, Ray Blanchard, Marc Breedlove, and Jan Kabatek for providing data, and Guillaume Martin for help with Mathematica. This work has been realized with the support of MESO@LR-Platform at the University of Montpellier. This is contribution 2022.050-SUD of the Institut des Sciences de l’Evolution de Montpellier (UMR CNRS 5554).

