## Supplementary materials for "Increased birth rank of homosexual males: disentangling the older brother effect and sexual antagonism hypothesis" (from another version of this preprint: version 4, 2022-11-26)

========================================

#### Supplementary materials

**Page**

**Appendix 1** ………………………. **2**
**Appendix 1b** ………………………. **4**
**Appendix 2** ………………………. **5**
**Appendix 3** ………………………. **5**
**Supplementary tables**
**Table S1** ………………. **7**
**Table S2** ………………. **8**
**Table S3** ………………. **9**
**Table S4** ………………. **10**
**Table S5** ………………. **11**
**Supplementary figures**
**Fig S1** ……………….. **12**
**Fig S2** ……………….. **13**
**Fig S3** ……………….. **14**
**References cited** ……………….. **15**

========================================

**Appendix 1. Relation between fertility and birth rank**

Consider a population of *N* families having fecundity $\lambda$ . The mean total number of children produced is $\lambda N$ .

<u>Probability of sampling a first-born</u>. The probability for a family to have no child is $e^{-\lambda}$ , so there are $N e^{-\lambda}$ families in such a case. The other $N(1-e^{-\lambda})$ families have at least one child, so there are $N(1-e^{\lambda})$ number of first born in the children population. The probability $R_1$ of sampling a firstborn, for a given $\lambda$ , is thus: $R_1(\lambda) = \frac{1-e^{-\lambda}}{\lambda}$ .

<u>Probability of sampling a second-born</u>. The probability for a family to have only one child is $e^{-\lambda}\lambda$ , so there are $N e^{-\lambda}\lambda$ families in such a case. The number of families with at least a second child is $N(1-e^{-\lambda}-\lambda e^{-\lambda})$ , and this is also the number of second born in the children population. The probability $R_2$ of sampling a second born is thus: $R_2(\lambda) = \frac{1-e^{-\lambda}(1+\lambda)}{\lambda}$

<u>Probability of sampling an individual with birth rank j</u>. The probability for a family of having fewer than *j* children is the sum of the probability of having *0, 1,.., j-1* children, thus $e^{-\lambda}\sum_{k=0}^{j-1}\frac{\lambda^k}{k!}$ . The number of families with at least *j* children is $N(1-e^{-\lambda}\sum_{k=0}^{j-1}\frac{\lambda^k}{k!})$ , and this is also the number of children of rank *j* in the population of children. The probability $R_j$ of sampling someone of rank *j* (with *j* ≥ 1) is thus:

$$R_j(\lambda)=\frac{1-e^{-\lambda}\sum_{k=0}^{j-1}\frac{\lambda^k}{k!}}{\lambda} \qquad \text{(A1.1)}$$

It is possible to check that $\sum_{j=1}^{\infty} R_j(\lambda)=1$ . Let's calculate $S_n=\sum_{j=1}^{n} R_j(\lambda)$ and then $\lim_{1}^{\infty} S_n$ .

$$S_n=\sum_{j=1}^{n} R_j(\lambda)=\frac{1}{\lambda}\sum_{j=1}^{n}(1-\sum_{0}^{j-1}P_j) \text{ , where } P_j=\frac{e^{-\lambda}\lambda^j}{j!}$$

$$\lambda S_n=(1-P_0)+(1-P_0-P_1)+(1-P_0-P_1-P_2)...+(1-P_0-P_1-..-P_{n-1})$$

$$\lambda S_n = n - nP_0 - (n-1)P_1 - (n-2)P_2 - \ldots - (n-(n-1))P_{n-1}$$

$$\lambda S_n = n - n\sum_0^{n-1} P_j + \sum_0^{n-1} j \cdot P_j$$

$\lim_1^\infty \lambda S_n = n - n\sum_0^\infty P_j + \sum_0^\infty j \cdot P_j = n - n + \lambda = \lambda$ , thus $\sum_{j=1}^\infty R_j(\lambda) = 1$

It is possible to check that $\overline{R(\lambda)} = \sum_{j=1}^\infty j \, . R_j(\lambda) = \frac{\lambda+2}{2}$ **.**

Let's calculate $M_n = \sum_{j=1}^n j \, . R_j(\lambda)$ and then $\lim_1^\infty M_n$ .

$$M_n = \sum_{j=1}^n j \, . R_j(\lambda) = \frac{1}{\lambda}\sum_{j=1}^n j(1 - \sum_0^{j-1} P_j) = \frac{1}{\lambda}(\frac{n(n+1)}{2} - \sum_{j=1}^n j\sum_0^{j-1} P_j)$$

$$\lambda M_n = \frac{n(n+1)}{2} - (P_0) - 2(P_0 + P_1) - 3(P_0 + P_1 + P_2) - \ldots - n(P_0 + P_1 + .. + P_{n-1})$$

$$\lambda M_n = \frac{n(n+1)}{2} - P_0\sum_{i=1}^n i - P_1\sum_{i=2}^n i - P_2\sum_{i=3}^n i - \ldots - P_j\sum_{i=j+1}^n i - .. - P_{n-1}\sum_{i=n}^n i$$

$$\lambda M_n = \frac{n(n+1)}{2} - \sum_{j=0}^{n-1} P_j \sum_{i=j+1}^n i$$

as $\sum_{j+1}^n i = \sum_1^n i - \sum_1^j i = \frac{n(n+1)}{2} - \frac{j(j+1)}{2}$

$$\lambda M_n = \frac{n(n+1)}{2} - \sum_{j=0}^{n-1} P_j(\frac{n(n+1)}{2} - \frac{j(j+1)}{2})$$

$$\lambda M_n = \frac{n(n+1)}{2}(1 - \sum_{j=0}^{n-1} P_j) + (\sum_{j=0}^{n-1} P_j \frac{j(j+1)}{2})$$

$$\lambda M_n = \frac{n(n+1)}{2}(1 - \sum_{j=0}^{n-1} P_j) + \frac{1}{2}(\sum_{j=0}^{n-1} j^2 . P_j + \sum_{j=0}^{n-1} j \, . P_j)$$

$$\lim_1^\infty \lambda M_n = \frac{1}{2}(E(P^2) + E(P)) = \frac{1}{2}(V(P) + E(P)^2 + E(P)) = \frac{\lambda + \lambda^2 + \lambda}{2}$$

where $V(P)$ and $E(P)$ are the variance and the mean, respectively, of a Poisson distribution.

Thus, $M_n = \frac{\lambda+2}{2}$ (A1.2)

**Appendix 1b. Sampling distribution of older sisters**

The probability distribution of older sisters is calculated considering that men with various male birth orders are sampled. The probability of observing *i* older sisters is the sum of the events of observing *i* sisters for all categories of male birth order (or number of older brothers):

$$prob(os=i)=\sum_{j=0}^{\infty} prob(os=i \wedge ob=j)$$

The probability of the joint event $os = i$ and $ob = j$ is the probability of sampling an individual with birth rank $i+j+1$, restricted to the number of corresponding cases. With $i+j$ older sibliongs, there are $2^{i+j}$ combinations, among which $C_{i+j}^{j}$ correspond to *i* sisters and *j* brothers. Thus,

$$prob(os=i \wedge ob=j)=R_{i+j+1}(\lambda)\, C_{i+j}^{j}\left(\frac{1}{2}\right)^{i+j}$$

and $$Prob(os=i)=\sum_{j=0}^{\infty} R_{i+j+1}(\lambda) C_{i+j}^{j}\left(\frac{1}{2}\right)^{i+j} \quad \text{(A1b.1)}$$

It is possible to verify that $\sum_{i=0}^{\infty}\sum_{j=0}^{\infty} R_{i+j+1}(\lambda) C_{i+j}^{j}\left(\frac{1}{2}\right)^{i+j}=1$ (details not shown). When sampling is random and there are no sex-biases for birth rank (e.g. absence of FBOE),

$prob(os=i)=prob(ob=i)$, thus Eq A1.1 and A1b.1 should be equivalent, i.e.

$$R_i(\lambda/2)=\frac{1-e^{-\lambda/2}\sum_{k=0}^{i-1}\frac{(\lambda/2)^k}{k!}}{\lambda/2}=\sum_{j=0}^{\infty} R_{i+j+1}(\lambda) C_{i+j}^{j}\left(\frac{1}{2}\right)^{i+j}$$

This equality is verified analytically (using Mathematica, not shown), and with modelling, for $\lambda$ values from 0.5 to 6 (not shown).

**Appendix 2. Unbiased estimate of fertility from sibling data**

For a population following a Poisson with parameter $\lambda$, or P($\lambda$), *n* mothers will generate $n\lambda$

children. Among these $n\lambda$ individuals, there are n.P($x$=$i$) families of size $i$, and i.n.P(x=$i$) individuals from a sib size of $i$. The probability of sampling an individual from a sib size of $i$ is thus : $i.n.P(x = i) / n\lambda$, or $i.P(x = i) /\lambda$.

Thus, the probability Pr(x = i) of sampling an individual from a family of $i$ siblings ($i$ >0), is :

$$Pr(x=i)=i\frac{P(x=i)}{\lambda}=i\lambda^i.\frac{e^{-\lambda}}{i!\lambda}=\lambda^{i-1}.\frac{e^{-\lambda}}{(i-1)!}$$

It is easy to show that : $\sum_{i=1}^{\infty} Pr(x=i)=\sum_{i=1}^{\infty} i\lambda^i.\frac{e^{-\lambda}}{i!\lambda}=\sum_{i=0}^{\infty}\lambda^i.\frac{e^{-\lambda}}{i!}=1$

The expectation is then :

$$m=\sum_{i=1}^{\infty} i.Pr(x=i)=\sum_{i=1}^{\infty} i.\lambda^{i-1}.\frac{e^{-\lambda}}{(i-1)!}=\sum_{i=1}^{\infty}(i-1)\lambda^{i-1}.\frac{e^{-\lambda}}{(i-1)!}+\sum_{i=1}^{\infty}\lambda^{i-1}.\frac{e^{-\lambda}}{(i-1)!}$$

Thus : $m=\sum_{i=0}^{\infty} i.\lambda^i.\frac{e^{-\lambda}}{i!}+\sum_{i=0}^{\infty}\lambda^i.\frac{e^{-\lambda}}{i!}=\lambda+1$

Or: $m-1=\lambda$ , suggesting that the mean number of brothers and sisters (not counting the sampled individuals) is an unbiased estimate of $\lambda$ (see Fig. S2). This is a particular case, for a Poisson distribution, of eq. 15.3.3 of Keyfitz & Caswell (2005).

**Appendix 3. Re-analysis of SBOE in Blanchard and Lippa (2021)**

To test for an SBOE, Blanchard and Lippa (2021) contrast groups of homosexuals and heterosexuals with only one sibling (thus total sibship of exactly 2): those with only one older sister (Group2 in their Table 1), and those with only one younger sibling (Group3 in their Table 1). This provides the following 2x2 contingency table:

| | Homo | Hetero |
|---|---|---|
| Group2 | 815 | 8134 |
| Group3 | 1482 | 16514 |

Rows and columns are not independent (Fisher exact test, P = 0.017, odds ratio = 1.12), and homosexual orientation increases when a younger sibling is replaced by an older sister (from Group3 to Group2): this is evidence, according to Blanchard and Lippa (2021), of an SBOE. However, the younger sib in Group3 is either a male or a female. Thus, both sex ratio and rank are changed between Groups 2 and 3. It would have been preferable to test Group2 vs Group3', where Group3' is the situation with only a younger sister, thus only the birth order change between Group2 and Group3', and not also the sex (between Group2 and Group3). A proxy of Group3' is Group3 divided by 2 (assuming a 1:1 sex-ratio). The corresponding Fisher exact test provides P = 0.039.

Blanchard and Lippa (2021) provide a replication in larger families, although, as they said, the groups used for the comparisons "*were approximately rather than exactly matched on family size.*" Thus, it cannot be considered that fertility effect is fully controlled for.

In conclusion, there is evidence for a SBOE in these data, with a marginal significance level (P-value = 0.04).

### Supplementary tables

**Table S1**. Functions used to model an older sibling effect. X= number of older brothers, or older sisters, for FBOE or SBOE, respectively. $p_0$ is the probability of a firstborn being homosexual. $\lambda$ is the mean fertility. The type of curve (family), and the corresponding parameters are indicated.

| Name | Form | Family | Parameters |
|---|---|---|---|
| f1 | $\frac{e^{a.X+b}}{1+e^{a.X+b}}.c+p_0$ | Logistic | $p_0$, a, b, c |
| f2 | $\frac{e^{a.X+b}}{1+e^{a.X+b}}.(1-p_0)+p_0$ | Logistic | $p_0$, a, b |
| f3 | $\frac{e^{a.X+b}}{1+e^{a.X+b}}$ , with $b=\log(\frac{p_0}{1-p_0})$ | Logistic | $p_0$, a |
| f4 | $a.X^{\gamma}+p_0$ | Geometric | $p_0$, a, $\gamma$ |
| f5 | $a.X+p_0$ | Linear | $p_0$, a |
| f6 | $p_0(for\ X=0), p_0+a_1+a.(X-1)(for\ X\geq 1)$ | Linear, with threshold | $p_0$, a, $a_1$ |
| f7 | $p_0(for\ X=0), p_0+a_1(for\ X>0)$ | Constant, with threshold | $p_0$, $a_1$ |
| f7’ | same as f7, with $a_1 = \mu.\lambda$ | Constant, with threshold | $p_0$, $\mu$ |
| f8 | $p_0+a_1.X+a_2.X^2$ | Polynomial | $p_0$, $a_1$, $a_2$ |

**Table S2**. Simulating the variation of mean birth rank in function of mean fertility, when FBOE, AE, or both, are present. FBOE was modelled by three possible functions describing an increase of probability of being homosexual with birth order (parameters $a$, $a_1$, and $\mu$, respectively), and AE was modelled an increase in $\lambda$ by a factor ($1+\beta$), conjointly with an increase in $p_0$ by a factor ($1+\alpha$). The parameter values for each simulation are given, as well as the resulting slope of the regression line of mean birth rank on mean fertility, standard error (SE) in parentheses, and the *P*-value of the test of the slope relatively to the expected value (0.5). Significant values ($P < 0.05$) are in bold.

| | Parameters | | | | | | |
|---|---|---|---|---|---|---|---|
| | FBOE | | | AE | | | |
| Effect | a | a1 | μ | α | β | Slope (SE) | *P*-value |
| AE | - | - | - | 1 | 1 | 0.50 ($<10^{-2}$) | 0.958 |
| AE | - | - | - | 2 | 1 | 0.50 ($<10^{-2}$) | 0.541 |
| FBOE (f5) | 0.2 | - | - | - | - | 0.67 (0.01) | **$< 10^{-3}$** |
| FBOE (f7) | - | 0.4 | - | - | - | 0.49 (0.01) | 0.975 |
| FBOE (f7' : f7 with a=μ.λ) | - | - | 0.1 | - | - | 0.53 (0.01) | **$< 10^{-3}$** |
| AE & FBOE | 0.2 | - | - | 2 | 1 | 0.53 ($<10^{-2}$) | **$< 10^{-3}$** |
| AE & FBOE | - | 0.2 | - | 2 | 1 | 0.51 ($<10^{-2}$) | **0.021** |
| AE & FBOE | - | 0.4 | - | 2 | 1 | 0.50 ($<10^{-2}$) | 0.544 |
| AE & FBOE | - | - | 0.1 | 2 | 1 | 0.53 ($<10^{-2}$) | **$< 10^{-3}$** |

**Table S3**. The relationship between mother's fertility and offspring sexual orientation, controlled for male birth rank and year of birth. The intercept of the logistic regression corresponds to the number of siblings (in logit units) for heterosexual men with a male birth order of 1 and an average year of birth. For the independent variables the estimate (β), standard error of the mean (SE), $X^2$, degrees of freedom (df), and the corresponding *P*-value are given for the three datasets (Indonesia, Greece, and France). Bold characters indicate significant ($P < 0.05$) effects.

| **Indonesia** | β (SE) | χ² (df) | P-value |
|---|---|---|---|
| Intercept | 0.90 (0.09) | - | - |
| Sexual orient. | -0.05 (0.09) | 0.3 (1) | 0.60 |
| Year of birth | -0.24 (0.04) | 35.3 (1) | **< $10^{-8}$** |
| Male birth order | - | 99.6 (6) | **< $10^{-8}$** |
| 2 | 0.42 (0.11) | - | - |
| 3 | 0.62 (0.13) | - | - |
| 4 | 1.12 (0.12) | - | - |
| 5 | 0.49 (0.26) | - | - |
| 7 | 1.18 (0.20) | - | - |
| 11 | 1.05 (0.31) | - | - |
| **Greece** | β (SE) | χ² (df) | P-value |
| Intercept | 0.16 (0.04) | | |
| Sexual orient. | 0.01 (0.07) | 0.01 (1) | 0.93 |
| Year of birth | -0.02 (0.03) | 0.28 (1) | 0.59 |
| Male birth order | | 153.2 (5) | **< $10^{-8}$** |
| 2 | 0.41 (0.07) | - | - |
| 3 | 0.92 (0.09) | - | - |
| 4 | 1.32 (0.20) | - | - |
| 5 | 1.91 (0.36) | - | - |
| 6 | 1.92 (0.36) | - | - |
| **France** | β (SE) | χ² (df) | P-value |
| Intercept | 0.31 (0.06) | - | - |
| Sexual orient. | -0.07 (0.07) | 1.0 (1) | 0.32 |
| Year of birth | -0.09 (0.03) | 6.8 (1) | **0.01** |
| Male birth order | | 71.3 (4) | **< $10^{-8}$** |
| 2 | 0.37 (0.08) | - | - |
| 3 | 0.83 (0.10) | - | - |
| 4 | 1.03 (0.31) | - | - |
| 5 | 0.95 (0.51) | - | - |

**Table S4**. The relationship between aunts' fertility and offspring sexual orientation, controlled for male birth rank and year of birth. The intercept of the regression corresponds to the number of cousins (in logit units for France) for heterosexual men with a male birth order of 1, an average year of birth, and an average number of aunts. For the independent variables the estimate (β), standard error of the mean (SE), $X^2$ or F, degrees of freedom (df), and the corresponding *P*-value are given for the two datasets (Indonesia, and France). Bold characters indicate significant ($P < 0.05$) effects.

| **Indonesia** | β (SE) | F (df) | *P*-value |
|---|---|---|---|
| Intercept | 4.80 (0.49) | - - | - |
| Sexual orient. | -0.13 (0.57) | 0.05 (1) | 0.82 |
| Year of birth | -0.21 (0.28) | 0.57 (1) | 0.45 |
| Male birth order | | 2.85 (6) | **0.01** |
| 2 | -0.24 (0.64) | - | - |
| 3 | 0.35 (0.89) | - | - |
| 4 | 0.06 (1.01) | - | - |
| 5 | 2.59 (2.08) | - | - |
| 7 | 2.33 (2.08) | - | - |
| 11 | 13.67 (3.57) | - | - |
| Number of aunts | 3.33 (0.26) | 158.9 (1) | **< $10^{-8}$** |

| **France** | β (SE) | $\chi^2$ (df) | *P*-value |
|---|---|---|---|
| Intercept | 0.51 (0.06) | - | - |
| Sexual orient. | 0.13 (0.07) | 3.16 (1) | 0.08 |
| Year of birth | -0.03 (0.03) | 0.73 (1) | 0.39 |
| Male birth order | | 3.61 (4) | 0.46 |
| 2 | -0.01 (0.08) | - | - |
| 3 | 0.15 (0.14) | - | - |
| 4 | 0.39 (0.36) | - | - |
| 5 | -13.3 (575) | - | - |
| Number of aunts | 0.70 (0.03) | 673 (1) | **< $10^{-8}$** |

**Table S5**. WAIC for the fit of the various functions describing an older sib effects. For each dataset and for each effect (OB: older brother, OS: older sister) the lowest WAIC is underlined. WAIC values lower than the minimum value plus 2 are in bold. The functions (f1 to f8) are defined in Table S1.

| | OB | | | OS | | |
|---|---|---|---|---|---|---|
| Function | Indonesia | Greece | France | Indonesia | Greece | France |
| f1 | 220.33 | 939.68 | **705.18** | 213.35 | 942.14 | **698.87** |
| f2 | 221.15 | 945.01 | **704.95** | 215.17 | 947.60 | **698.72** |
| f3 | **218.02** | **937.54** | **705.73** | **212.18** | 942.76 | **700.12** |
| f4 | **218.14** | **938.15** | **705.83** | **211.56** | **941.62** | **700.08** |
| f5 | **217.90** | **937.37** | **705.67** | **212.32** | 942.42 | **700.09** |
| f6 | **218.78** | **938.98** | **705.89** | **212.29** | **941.24** | 701.01 |
| f7 | **217.35** | **937.40** | **704.28** | **210.50** | **939.95** | **699.66** |
| f8 | 219.75 | **939.04** | 706.27 | **212.32** | **941.23** | 701.73 |

### Supplementary figures

**Figure S1**. Simulating the variation in mean birth order as a function of mean fecundity. 2000 families were generated, with fecundity from a Poisson distribution of parameter λ. The birth order of all siblings was recorded and the mean birth order from a random sample of 300 individuals was computed. This was replicated 50 times for each λ from 0.5 to 6. The dotted line represents the theoretical relationships (mean rank = λ/2 +1, see results)

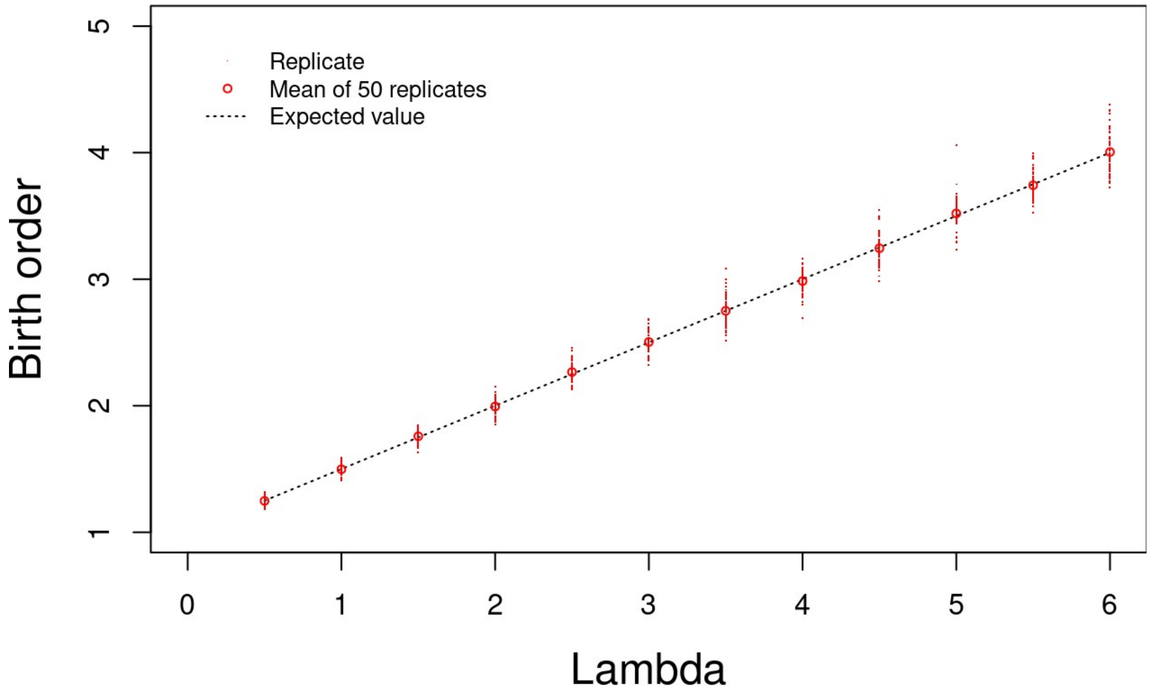

**Figure S2**. Comparison of various estimates of fecundity from sibling data. 2000 families were generated with family fecundity drawn from a Poisson distribution of parameter λ. For each offspring individual the full fertility of his mother was recorded and, from a random sample of 300 individuals, the mean fertility λ of the population was estimated using 4 methods. This was replicated 50 times for each λ from 0.5 to 6. The four methods are: 1) mean of full sibling size, with full sibling size = brothers+sisters+sampled individual ("Without correction"), 2) mean of full sibling size corrected with a zero-truncated Poisson distribution ("Correction bias 1"), 3) mean of full sibling size corrected for sibling size ("Correction bias 2"), as proposed by Berglin (1980), and 4) mean of number of brothers and sisters, without counting the sampled individual ("Correction biases 1 and 2"). The dotted line represents the expected λ values.

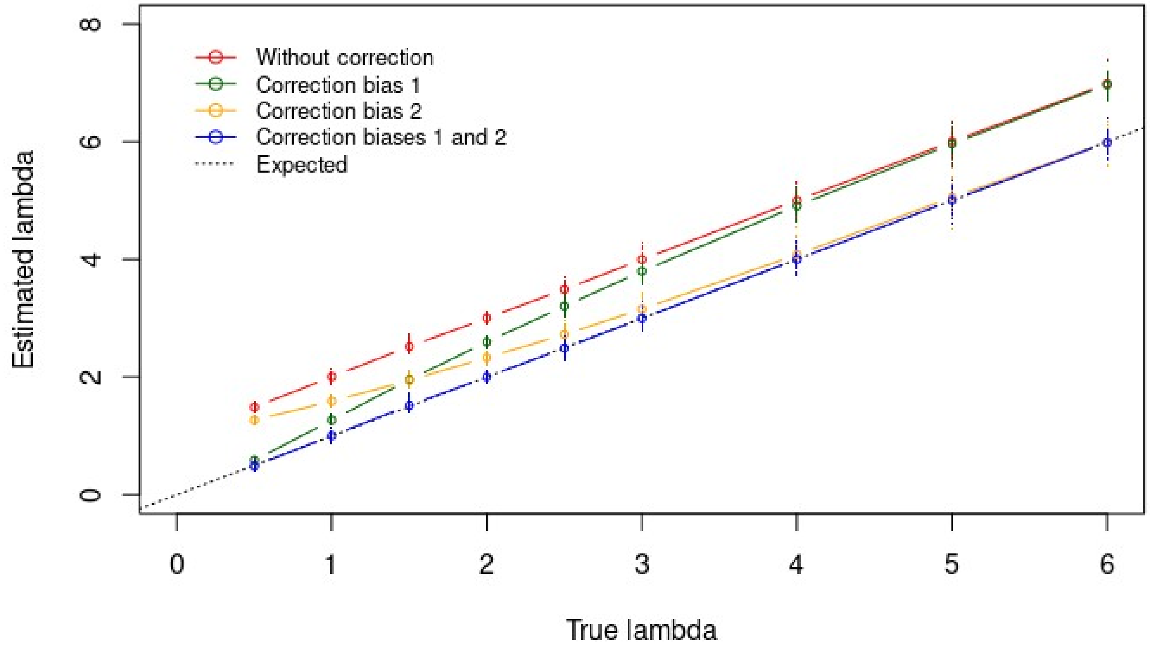

**Figure S3**. Simulating the effect of FBOE and/or AE on the relationship between mean birth rank and mean fertility. 6000 families were generated, with family fecundity drawn from a Poisson distribution of parameter λ. AE was absent (left panels A, C, and E), or present for ¼ of the families (right panels B, D, and F). When present it was modeled with parameters $\alpha = 2$ and $\beta = 1$. FBOE was absent (row 1, panels A and B), or present (rows 2 and 3). When present it was generated with function f5 with a = 0.2 (row 2, panels C and D), or function f7' with $\mu = 0.05$ (row 3, panels E and F). In all case, $p_0 = 0.05$. From a random sample of 200 homosexual and 200 heterosexual men, the mean male birth rank and the mean fecundity for males was computed for each sex orientation (red: heterosexual men; blue: homosexual men). This was replicated 200 times (dots) for each λ from 2 to 8 (thus from 1 to 4 considering the resulting fecundity for males). The dotted line represents the theoretical relationships (mean rank = λ/2 +1, see results). Data above 4 on the x-axis or y-axis are not represented.

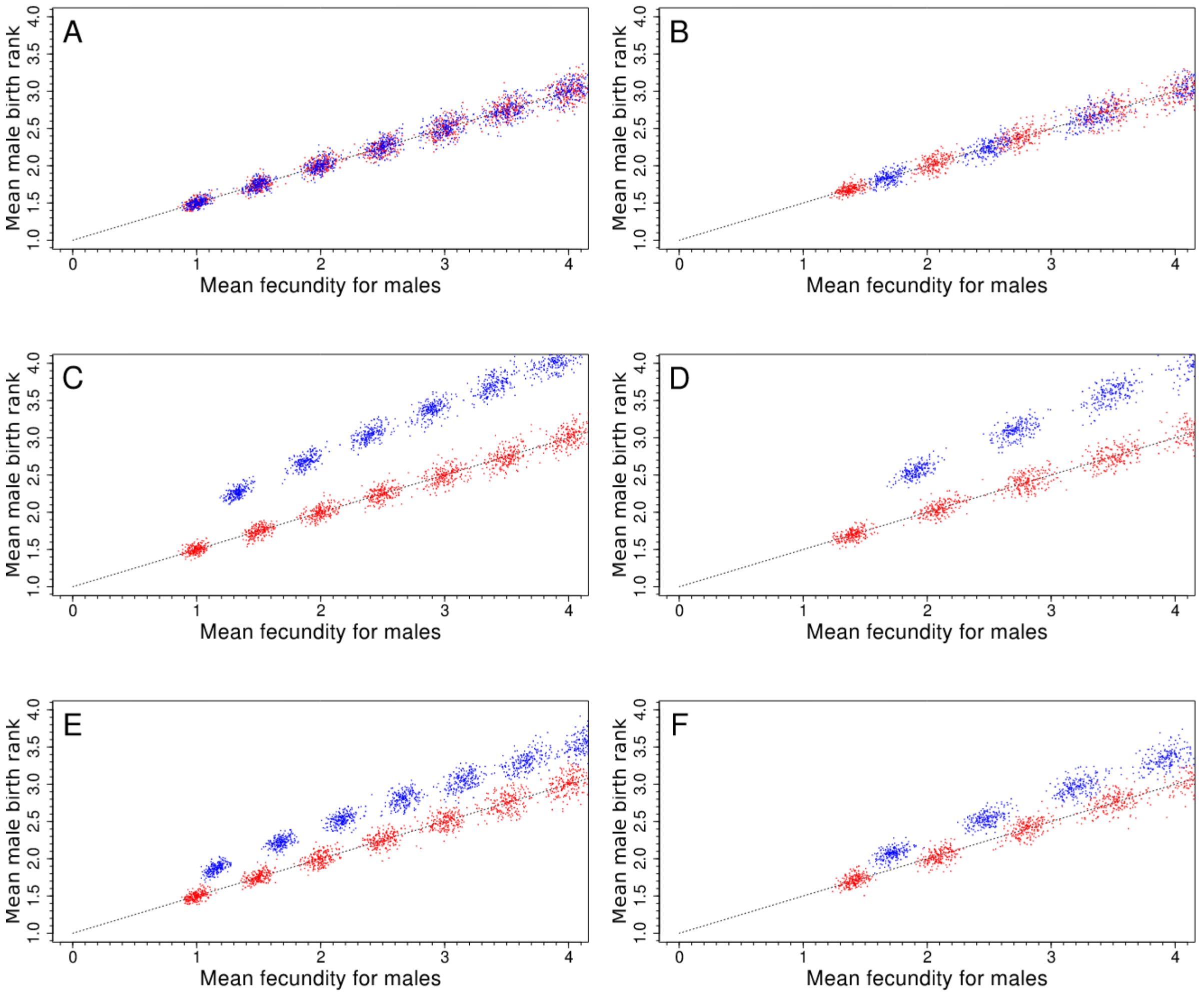
